# The rENM Framework: A Modular System for Reconstructing and Analyzing Long-Term Ecological Niche Dynamics

**DOI:** 10.64898/2026.08.06.741224

**Authors:** John L. Schnase, Mark L. Carroll, Paul M. Montesano, Virginia A. Seamster

## Abstract

Retrospective ecological niche modeling (rENM) combines historical species occurrence records with historical environmental data to reconstruct the spatio-temporal dynamics of species-environment relationships under changing conditions. Despite growing recognition that those relationships can be nonstationary, time-series approaches to ecological niche modeling remain uncommon, and the tools to support them at scale are limited. Here, we describe the rENM Framework, an experimental, open-source suite of R packages that automates a complete rENM workflow spanning data preparation, ensemble time-series construction, trend analysis, AI interpretation, and report generation. The framework integrates eBird occurrence records with environmental variables derived from NASA’s MERRA-2 reanalysis across a 45-year study period (1980–2024) and executes a complete analysis for any species with eBird data through a single function call. By treating climatic suitability as a dynamic ecological response surface rather than a static baseline, the framework produces the following analytical products that complement conventional ecological niche modeling approaches: suitability time series, long-term trend and acceleration maps, centroid displacement estimates, bioclimatic velocity metrics, variable contribution trajectories, and hotspot analyses identifying areas of accelerating suitability decline. We illustrate the framework’s outputs with a representative run for Cassin’s Sparrow (*Peucaea cassinii*), a grassland species of conservation concern in the arid southwestern United States and the focal species throughout our development work. The framework’s automated, unsupervised pipeline makes systematic application across large numbers of species tractable, with direct implications for conservation assessments, such as State Wildlife Action Plans, where species-specific analytical capacity is often limited by available resources. The rENM Framework is openly available on GitHub and archived on Zenodo.

## Introduction

Retrospective ecological niche modeling (rENM) combines historical species occurrence records with historical environmental data to reconstruct and analyze the spatio-temporal dynamics of species-environment relationships under changing environmental conditions [1,2]. The rENM Framework is a modular suite of R packages designed to support rENM analyses. By reconstructing long-term niche dynamics, rENMs provide an empirical basis for addressing biological questions and informing conservation assessments. Rather than treating ecological niche models (ENMs) as static representations, the rENM Framework uses time-structured modeling to reveal long-term trends in environmental suitability for a species, including acceleration or deceleration of those trends, directional change in suitability, decadal changes in environmental structure, bioclimatic velocity, and hotspots of potential climate change vulnerability.

Although this analytical approach is broadly applicable across taxa, the current implementation of the rENM Framework focuses on North American bird species and climatic suitability. Birds provide a strategic starting point because of the temporal depth, geographic coverage, and data quantity and quality afforded by eBird, the globally curated, continuously updated citizen-science archive of bird observations maintained by the Cornell Lab of Ornithology [3]. The framework spans 45 years (1980–2024) and integrates eBird occurrence records with environmental variables derived from the National Aeronautics and Space Administration’s (NASA’s) Modern-Era Retrospective Analysis for Research and Applications, Version 2 (MERRA-2) reanalysis [4,5]. The approach has shown promise in our initial studies, producing biologically interpretable results and yielding new insights into the ecology of the species we have studied; however, it remains experimental, and its general applicability has yet to be established. With this paper, we make the rENM Framework publicly available and invite the broader community to evaluate, replicate, and extend the approach.

### ENM background and limitations

Ecological niche modeling (ENM), also referred to as species distribution modeling (SDM), consists of a set of techniques and tools that use species occurrence records and environmental data to map the relative suitability of habitats [6,7]. The approach is grounded in Hutchinson’s formulation of the niche as an *n*-dimensional hypervolume [8] and operationalizes the distinction between the fundamental niche, the full range of abiotic conditions permitting persistence, and the realized niche, the subset of that space actually occupied given biotic interactions, dispersal history, landscape connectivity, and other factors [9]. ENM is used across a wide range of disciplines, including fields as diverse as biogeography and phylogeny [10], epidemiology [11], invasion biology [12], and archaeology [13]. In recent years, ENMs have become particularly important in understanding the influence of climate change on the geographic distribution and conservation status of species [14–17].

Despite its broad utility, ENM carries well-documented limitations. Among the most important, ENM workflows often rely on environmental predictors summarized as multi-decadal averages. Because these predictors are not always temporally aligned with the occurrence records used in model development, they can mask temporal variation in the conditions associated with species occurrence [18,19]. A deeper problem arises when species–environment relationships themselves change through time, a phenomenon known as nonstationarity [20,21]. With nonstationarity, the statistical properties of an ecological process, such as the relationships among climatic drivers and species occurrence, vary across space or time rather than remaining constant [20,21]. When nonstationarity is present, a single time-averaged predictor surface may inadequately represent the factors influencing species distributions, not merely because it smooths over variation, but because the underlying relationships being modeled may themselves be shifting [1,2,20,21].

This limitation has important consequences in two contexts. First, time-averaged representations are poorly suited for highly mobile or behaviorally responsive species whose distributions track changing environmental conditions across seasonal and interannual time scales [1,2,22–24]. This includes important classes of birds, such as grassland species that respond to rainfall, vegetation productivity, and drought cycles, as well as nomadic, irruptive, and migratory species whose distributions track shifting resources from season to season and year to year [25,26].

Second, time-averaged predictors complicate forward projection. Conventional forecasts typically calibrate models against a single present-day baseline and then project those relationships into future climate scenarios. When species–environment relationships change through time, however, a single baseline may provide an incomplete representation of the processes governing distributional change. This is particularly consequential in conservation applications where the goal is not merely to characterize current habitat distribution but to understand how distributions have changed through time and to use that knowledge to inform policy and management decisions [27–30]. Historical trajectories of suitability provide important context for that understanding, offering a more empirically grounded departure point for forecasting than any single baseline snapshot can [28,29].

Addressing nonstationarity in ecological and distribution models is an active area of methodological development. Current approaches include generalized additive models for capturing nonlinear predictor-response relationships over time, dynamic linear models in which regression coefficients are estimated as random walks, and spatially varying coefficient models that allow the relationships between predictors and responses to differ across geographic space [20]. More recently, spatiotemporal models have been extended to allow the variance structure of spatial processes to change through time, enabling detection of trends in the patchiness or spatial heterogeneity of population distributions as environmental conditions shift [21]. These methods share a common conceptual logic: rather than assuming that the relationships governing a species’ distribution are fixed and transferable, they treat those relationships as quantities that may need to be estimated and tracked as conditions evolve. The rENM Framework operates in this same spirit and complements these approaches by focusing on the temporal dynamics of climatic suitability reconstructed from historical occurrence and environmental records.

### Project motivation and history

Our work is motivated by the need to characterize the spatial and temporal dynamics of species’ relationships to changing environments. Despite growing recognition of nonstationarity as a core challenge for ENM, retrospective time-series approaches remain far less common than their potential warrants, and the tools to support them at scale are limited [19,29]. One reason is that these approaches can require the construction of complex, multi-member time series of ENMs spanning months, years, or decades. The rENM Framework is intended as an enabling technology that addresses this unmet need. In the rENM Framework, climatic suitability is treated not as a static representation of a species’ niche but as a dynamic ecological response surface that changes as environmental conditions shift across decades. The evolving suitability surface itself becomes an object of study, one that the framework can reconstruct automatically, offering a practical means of advancing work in this area [17,31–33].

The rENM Framework grew out of an ongoing series of retrospective analyses of Cassin’s Sparrow (*Peucaea cassinii*), a ground-dwelling grassland species of conservation concern in the arid southwestern United States and northern Mexico [1,2,34–37]. Those studies revealed that modeled estimates of climatic suitability for the species vary substantially depending on the temporal frame, spatial extent, and environmental predictors considered, and that variability appears to mirror the inconsistent picture of the species’ conservation status as reported across the literature [1,2,34]. Furthermore, retrospective analysis using temporally aligned occurrence and environmental data, with predictor variables independently selected for each time interval, revealed long-term directional shifts and regionally divergent trends in suitability that conventional temporally static ENM approaches could not [1]. The rENM Framework codifies the primary analytical workflow developed in these studies. We release it here as an open research platform in the hope that community engagement will advance the tools themselves and the science they are designed to support.

## Materials and Methods

The rENM Framework comprises a modular suite of seven R packages organized around the sequential steps of the rENM analytical workflow (Fig 1). A core utilities package (rENM.core) provides shared infrastructure used across all other modules. A top-level orchestration package (rENM) coordinates execution of the complete pipeline, allowing the entire workflow to be run for a target species with a single function call. The remaining five packages correspond to the major functional stages of the analysis: data assembly (rENM.data), time-series construction and modeling (rENM.model), time-series analysis (rENM.analysis), generative AI (GenAI) interpretation (rENM.ai), and report generation (rENM.reports).

**Fig 1.**
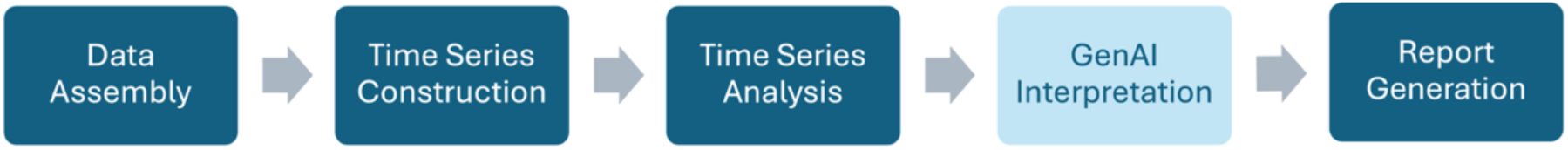
rENM processing pipeline. Figure shows the main processing steps in the rENM workflow. The GenAI Interpretation stage is optional and experimental.

This section summarizes each package’s architecture and primary functions at a high level. Algorithmic detail and function-specific inputs and outputs are covered in the User Manual [38] and package reference manuals. The framework is available on GitHub [39] and archived on Zenodo [40]. Installation instructions, technical reference manuals, workflow documentation, related publications, and access to example input datasets and analysis outputs are provided through the project’s online documentation (https://github.com/rENM-Framework).

### Host environment

The rENM Framework is implemented in R (version 4.1.0 or later) [41]; development and testing were done using the RStudio integrated development environment [42]. Framework packages are installed directly from the framework’s GitHub repository [39] or the Zenodo archive [40]. The framework is designed to operate on standard desktop or laptop computing environments and relies primarily on widely used open-source R packages and geospatial analysis libraries. Two external open-source software dependencies, LibreOffice and Chromium, are required for document rendering and report generation. The system has been developed and tested on macOS and is expected to be compatible with Linux environments. Because some of the framework’s parallel processing mechanisms are not consistently available on Windows, computationally intensive operations may execute sequentially, resulting in longer run times.

The framework relies on a simple filesystem structure. At the top level is the user’s `project` directory containing a `data` directory that stores input datasets, including species occurrence records and environmental variables; model outputs and analysis results are written to a species-specific `runs` directory (Fig 2).

**Fig 2.**
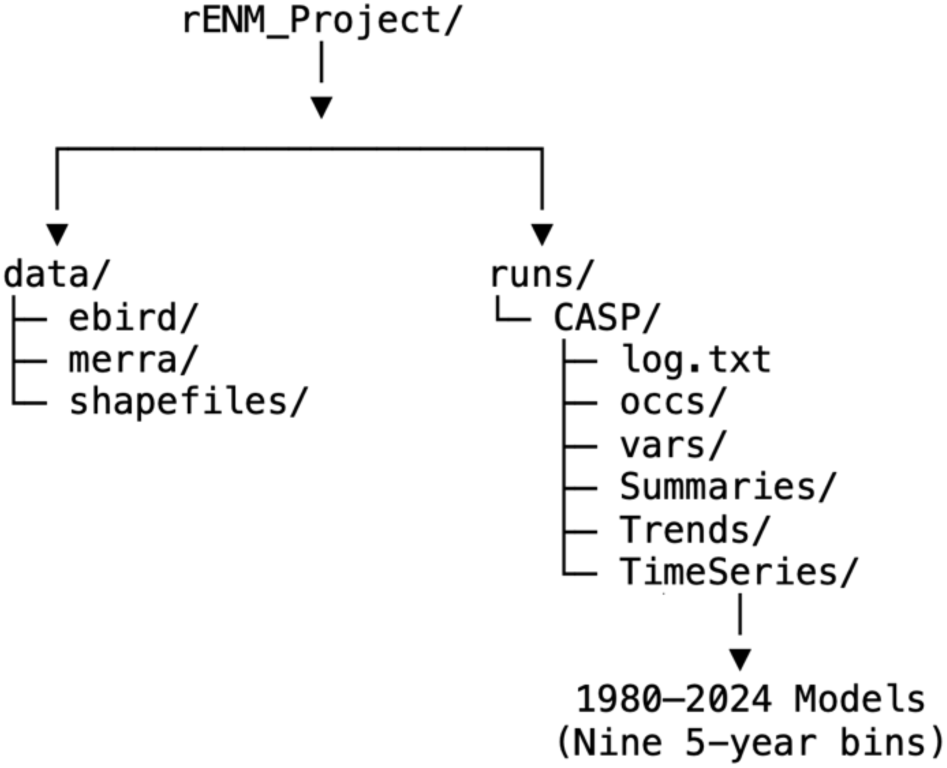
rENM project directory structure. Figure shows the layout of the major directories operated on by the rENM Framework’s processing workflow.

The current release of the rENM Framework supports two categories of model input data, both of which must be provisioned by the user in the host environment’s `data` directory before analysis can begin. For species occurrence records, the framework accommodates downloaded eBird Basic Dataset (EBD) files [43]. For environmental predictors, the framework supports gridded variables derived from NASA’s MERRA-2 dataset, a global atmospheric reanalysis that combines past observations with numerical models to generate a consistent, multi-decadal time series of physical drivers of the Earth system. Important for the work described here, MERRA-2 extends from 1979 to the present, and the temporal resolution of its variables is hourly [4,5]. The framework uses two types of MERRA-2-derived predictors: microclimatic and ecosystem functional variables drawn directly from MERRA-2 output and bioclimatic variables modeled after WorldClim predictors but derived from MERRA-2 temperature and precipitation fields. We refer to the latter as MERRAclim-2 variables [1,35,44] (S1 Appendix. MERRA-2 and MERRAclim-2 variables.). The framework uses United States Geological Survey (USGS) National Gap Analysis Program (GAP) species range maps [45] in several of its output products; these also must be provisioned by the user prior to an analysis run. The details on accessing required inputs can be found in the Data Availability Statement below.

### Framework modules

The following sections provide a high-level description of the major capabilities of each framework module and its role within the rENM analytical workflow. Detailed technical documentation, including function reference manuals, algorithm descriptions, parameter specifications, implementation notes, and usage examples, is available in the GitHub repositories associated with each package and in the R help pages accessible from within RStudio.

### rENM.core

The rENM.core package provides the foundational utilities used throughout an rENM analysis (Table 1). The package is intentionally lightweight, focused on project organization, metadata discovery, and reproducibility. Its central function, rENM_project_dir(), resolves the active project root from the user’s environment, thereby ensuring that downstream packages locate the user’s project files consistently regardless of where the framework software is installed or run. All other rENM packages depend on the functions this package provides.

**Table 1.** rENM.core package functions.

| Function | Description |
| --- | --- |
| rENM_project_dir() | Resolve the active rENM project root |
| get_species_info() | Retrieve standardized metadata for a species by alpha code |
| show_species() | Inspect the catalog of provisioned species |
| show_variables() | Inspect the catalog of provisioned environmental variables |

### rENM.data

The rENM.data package provides tools to prepare inputs to the rENM analysis workflow (Table 2). Processing proceeds in three steps. In the first step, occurrence records are extracted from downloaded eBird EBD files, filtered for valid geographic coordinates, and aggregated into nine five-year temporal bins representing the 45-year study period supported by the framework. In the current implementation, these intervals span 1980–2024. Duplicate records are then removed, and an optional spatial thinning step can be applied to reduce sampling bias. The thinning algorithm enforces a minimum distance between retained points that can be set by the user. Two thinning implementations are available: thin_occurrences(), which processes bins sequentially, and thin_occurrences2(), a parallel implementation that reduces runtime on multicore systems. An optional record cap can also be applied to limit excessively high record densities within any bin.

**Table 2.**
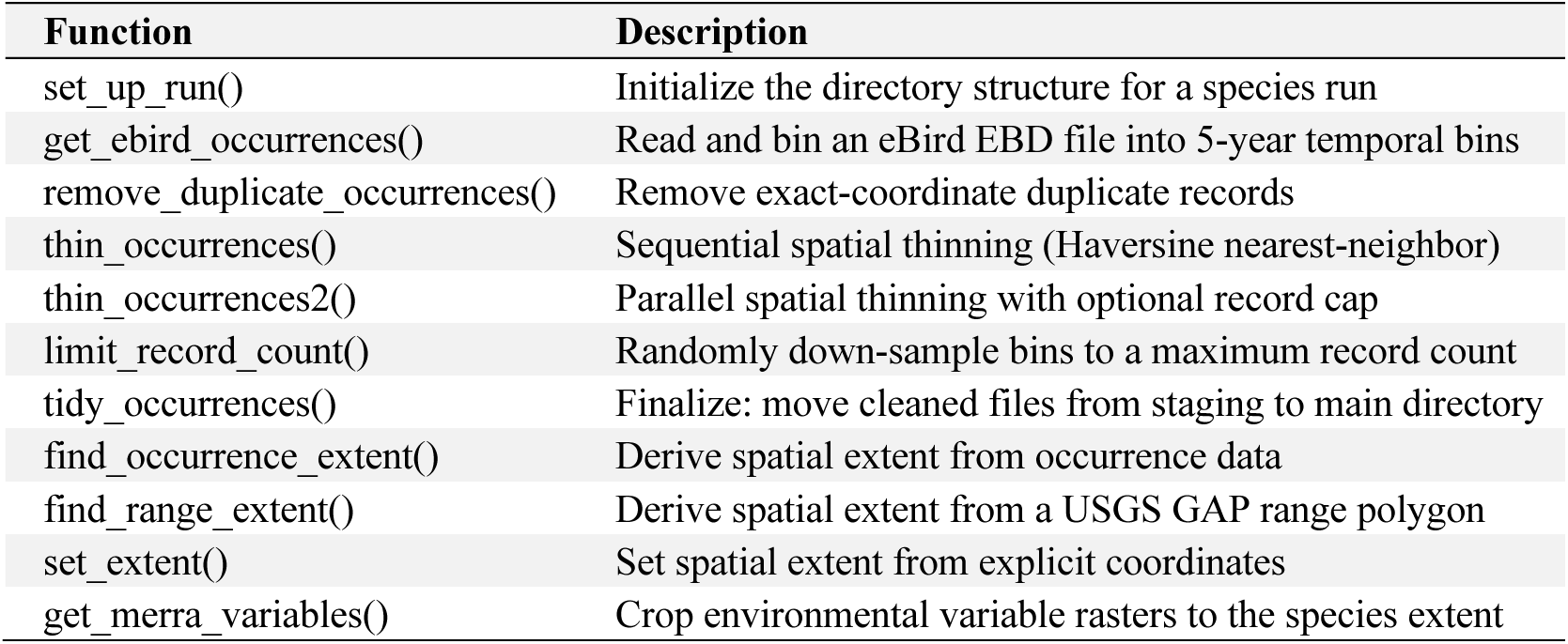
rENM.data package functions.

| Function | Description |
| --- | --- |
| `set_up_run()` | Initialize the directory structure for a species run |
| `get_ebird_occurrences()` | Read and bin an eBird EBD file into 5-year temporal bins |
| `remove_duplicate_occurrences()` | Remove exact-coordinate duplicate records |
| `thin_occurrences()` | Sequential spatial thinning (Haversine nearest-neighbor) |
| `thin_occurrences2()` | Parallel spatial thinning with optional record cap |
| `limit_record_count()` | Randomly down-sample bins to a maximum record count |
| `tidy_occurrences()` | Finalize: move cleaned files from staging to main directory |
| `find_occurrence_extent()` | Derive spatial extent from occurrence data |
| `find_range_extent()` | Derive spatial extent from a USGS GAP range polygon |
| `set_extent()` | Set spatial extent from explicit coordinates |
| `get_merra_variables()` | Crop environmental variable rasters to the species extent |

In the next step, the spatial extent of the analysis is defined. Three options are available for this: find_occurrence_extent() derives the extent from the species’ occurrence records using a centered percentile bounding box; find_range_extent() derives it from a USGS GAP range polygon with optional symmetric padding [45]; and set_extent() accepts explicit bounding box coordinates. Finally, get_merra_variables() copies MERRA-2 and MERRAclim-2 predictor rasters from the user’s base `data` directory collection into the `run` directory, cropping each layer to the defined spatial extent. The user has the option of moving all available environmental variables from the base `data` collection or selecting a subset. The resulting occurrence files and cropped predictor rasters constitute the complete set of time series inputs expected by rENM.model in the next processing step of the workflow. These data preparation steps store intermediate products in temporary system directories and in the `run` directory’s `_occ` and `_vars` directories before being staged into the appropriate temporal bins within the `TimeSeries` directory (Fig 2).

### rENM.model

The rENM.model package implements the modeling and time-series reconstruction components of the rENM Framework (Table 3). It transforms the occurrence records and environmental variables prepared by rENM.data into ensemble ENMs constructed from temporally aligned occurrence and environmental data for each five-year bin. The result is a time series of climatic suitability estimates that span the past 45 years.

**Table 3.** rENM.model package functions.

| Function | Description |
| --- | --- |
| stage_occurrences() | Copy occurrence files into TimeSeries bins |
| stage_all_variables() | Copy predictor rasters into TimeSeries bins |
| screen_by_convergence1() | Convergence-based variable screening via MaxEnt (requires Java) |
| screen_by_convergence2() | Convergence-based variable screening via maxnet (no Java dependency) |
| reduce_covariance() | Remove collinear predictors via adaptive VIF screening |
| stage_screened_variables() | Copy screened predictors into TimeSeries bins |
| create_ensemble_model() | Fit an ensemble ENM for a single species and time bin |
| create_timeseries() | Run ensemble models across all time bins in parallel |
| create_range_map() | Produce a binary presence-absence range map |
| plot_suitability() | Plot a continuous climatic suitability raster |
| save_suitability_plot() | Save a suitability plot to file |
| rank_variable_importance() | Parse and rank variable importance from an SDM model report |

The modeling process begins by staging occurrence records into the `TimeSeries` directory where they are organized into nine subdirectories corresponding to the five-year temporal bins of the analysis. Environmental variables are then staged to the corresponding bins to ensure temporal alignment. Users may either stage the complete collection of MERRA-2 and MERRAclim-2 variables assembled during the rENM.data preparation step or first apply an optional Monte Carlo–based screening procedure to identify the highest-contributing predictors for each five-year interval.

This screening option addresses two challenges that have historically complicated the use of large environmental datasets, particularly those derived from global climate models [35–37]. First, the size, complexity, and novelty of these datasets make variable selection difficult. For example, MERRA-2 contains more than 600 modeled variables, including radiation fluxes, soil moisture estimates, cloud properties, and other environmental measures that are rarely used in conventional ENM workflows. The framework’s screening procedure provides a computationally tractable means of evaluating the utility of these expanded variable collections.

Second, the screening procedure is biologically informed. It adapts the bivariate ensemble strategy introduced by Schnase and Carroll [35], in which large numbers of independent ENM runs, each operating on a randomly selected pair of predictors, converge on an estimate of the most influential variables within the full predictor set. Screening is performed independently for each five-year temporal bin, thus allowing convergence to be influenced only by the species-specific occurrence records and temporally aligned environmental variables within that interval. Permutation importance is calculated for each predictor pair; convergence is achieved when both the mean permutation importance of the top-*n* variables and the membership of the top-*n* set stabilize across successive iterations. The number of predictory-pair evaluations needed to achieve convergence will vary depending on occurrence record density and random sampling effects. The value of *n* may be specified by the user or determined adaptively based on the size of the predictor collection or occurrence record density. Variables that consistently emerge as top contributors are subsequently staged for time-series construction.

Two implementations of the screening procedure are available. The first function, screen_by_convergence1(), uses Java-based MaxEnt through the dismo R package [46]; screen_by_convergence2() uses maxnet and has no Java dependency [47], making it the recommended option for most users. Both implementations support parallel processing and can use available computational resources on the host system to accelerate screening. An optional covariance-reduction step may then be applied to remove collinear predictors from the staged variable sets using adaptive variance inflation factor (VIF) screening [48].

The create_timeseries() function makes parallel calls to create_ensemble_model() across all temporal bins to build the multi-decadal suitability time series that becomes the input to downstream analyses. The create_ensemble_model() function fits an ensemble of five ENMs for each five-year bin. The five algorithms include maxnet, random forest (RF), boosted regression trees (BRT), generalized linear models (GLM), and multivariate adaptive regression splines (MARS). These methods were selected because each is well-suited to presence-only data of the kind provided by eBird. In addition, this set spans a range of linear, nonlinear, and machine learning approaches that capture complementary aspects of the species-environment relationship [49,50]. Ensemble results are computed as an unweighted average across the five algorithms, a strategy that reduces sensitivity of suitability estimates to the idiosyncrasies of any single method and has been shown to improve model performance and transferability relative to single-algorithm models [51]. All five algorithms are accessed through the sdm R package using its default parameterizations [52].

### rENM.analysis

The rENM.analysis package produces the principal analytical outputs of the rENM Framework (Table 4). It transforms the suitability surfaces generated by rENM.model into five classes of analytical products: temporal trends, spatial extent summaries, centroid and bioclimatic velocity metrics, hotspot assessments, and variable contribution analyses. Temporal trend analyses use Theil-Sen median slope estimation and Mann-Kendall statistics to quantify changes in climatic suitability through time and identify areas of accelerating or decelerating change (i.e., suitability change trend) [53,54]. Spatial extent analyses summarize the distribution of positive, negative, and neutral suitability trends across the modeled landscape, within the species’ USGS GAP range, and at the state level. Centroid analyses quantify shifts in the suitability-weighted geographic center of the species’ climatic niche, including estimates of direction, distance, and velocity. Hotspot analyses identify regions exhibiting increasingly rapid declines in suitability that may warrant further investigation as areas of potential climate vulnerability. Finally, variable contribution analyses identify the environmental predictors most strongly associated with suitability in the time series models and evaluate how their relative importance changes through time. The outputs of these analyses are subsequently consumed by rENM.ai and rENM.reports.

**Table 4.** rENM.analysis package functions.

| Function | Description |
| --- | --- |
| find_suitability_trend() | Compute per-cell Theil-Sen trends and Mann-Kendall statistics |
| find_suitability_change_trend() | Compute trends in suitability rate of change |
| find_trend_percentages() | Summarize positive, negative, and zero trend proportions across modeled extent |
| find_range_change_percentages() | Summarize trend proportions within the USGS GAP range |
| create_state_trend_analysis() | Per-state suitability trend map and statistics |
| find_weighted_centroid() | Compute suitability-weighted spatial centroids |
| analyze_weighted_centroids() | Fit Bayesian trends to centroid latitude and longitude |
| find_bioclimatic_velocity() | Estimate climate-space displacement direction, distance, and velocity |
| find_hot_spots() | Identify cells with accelerating suitability declines |
| create_hot_spot_map() | Map hotspot regions within states intersecting the GAP range |
| create_suitability_change_map() | Full workflow for suitability change trend visualization |
| gather_variable_contributions() | Consolidate per-year variable importance files |
| summarize_variable_contributions() | Frequentist and Bayesian trend analysis of variable contributions |
| plot_trend() | Plot a climatic suitability trend raster |
| plot_trend_with_centroids() | Plot suitability trend with centroid shift overlay |
| plot_suitability_change_trend() | Plot suitability change trend raster |
| save_trend_plot() | Save a suitability trend map to file |
| save_trend_plot_with_centroids() | Save a suitability trend map with centroid displacement vector to file |

### rENM.ai

The rENM.ai package integrates artificial intelligence (AI) into the rENM workflow as an optional, experimental interpretive step (Table 5). Its operation proceeds in three steps. First, assemble_ai_package() collects key outputs produced by the workflow up to this point, such as suitability trend rasters, summary statistics related to suitability trend proportions, variable contribution tables, centroid displacement metrics, and species metadata, and organizes them into a compressed data bundle. A detailed analytical prompt is assembled alongside these data that will provide the destination AI system with the contextual information needed to analyze and summarize the rENM workflow’s outputs. The assembled package is then submitted to one of two large language models (LLMs), ChatGPT or Claude, via submit_to_chatgpt() or submit_to_claude(). The framework uploads the information through the OpenAI or Anthropic API and retrieves a Microsoft Word (.docx) report in return. The choice of provider affects response characteristics, processing time, and per-run cost. Finally, render_ai_docx() converts the returned .docx report to .pdf via LibreOffice for inclusion in a framework-generated analysis rENM report. Given the experimental nature of these AI outputs, they are intended as preliminary narratives to support expert review and should not be treated as validated ecological interpretations.

**Table 5.** rENM.ai package functions.

| Function | Description |
| --- | --- |
| `assemble_ai_package()` | Collect rENM outputs and assemble AI-ready data bundle with prompt |
| `submit_to_chatgpt()` | Submit data bundle to OpenAI and retrieve .docx report |
| `submit_to_claude()` | Submit data bundle to Anthropic and retrieve .docx report |
| `submit_to_claude_diag()` | Diagnose a failed `submit_to_claude()` response |
| `render_ai_docx()` | Convert AI-generated .docx report to .pdf via LibreOffice |

### rENM.reports

The rENM.reports package is the final stage of the rENM pipeline (Table 6). The package provides functions that consume the trend results, spatial metrics, summary statistics, map products, and AI analyses generated by previous steps in the workflow and organizes them into a coherent, paginated document. Report generation proceeds in three steps. First, gather functions collect and composite the raw outputs of the analysis pipeline into map contact sheets, merged statistics files, and variable trend composites. Table and page assembly functions then transform those intermediate products into formatted summary tables and single-page .pdf layouts covering suitability trends, centroid movement, state-level analyses, variable contributions, and time-series overviews. Finally, assemble_final_report() combines the individual pages into a single paginated .pdf document.

**Table 6.** rENM.reports package functions.

| Function | Description |
| --- | --- |
| `gather_suitability_maps()` | Assemble suitability map contact sheet |
| `gather_range_maps()` | Assemble binary range map contact sheet |
| `gather_suitability_trend_stats()` | Merge state-level suitability and hotspot statistics |
| `gather_top_variable_trend_maps()` | Assemble variable trend map composites |
| `create_suitability_trend_summary_table()` | State-level GAP range and hotspot summary table |
| `create_centroid_trend_summary_table()` | Centroid shift and regression summary table |
| `create_variable_trend_summary_table()` | Variable trend statistics summary table |
| `assemble_suitability_trends_page()` | Single-page .pdf: suitability trend and change-trend maps |
| `assemble_centroid_trends_page()` | Single-page .pdf: centroid trend map and table |
| `assemble_state_trends_page()` | Single-page .pdf: state trend map, hotspot map, and table |
| `assemble_variable_trends_page()` | Single-page .pdf: variable contributions plot and table |
| `assemble_variable_trend_maps_page()` | Single-page .pdf: variable trend map composites |
| `assemble_suitability_timeseries_page()` | Single-page .pdf: suitability time series contact sheet |
| `assemble_range_timeseries_page()` | Single-page .pdf: binary range time series contact sheet |
| `assemble_final_report()` | Combine all pages into a single paginated .pdf report |

### rENM

The rENM package is the top-level orchestration layer of the rENM Framework (Table 7). It provides a single exported function, rENM(), which automatically executes the complete analytical pipeline for a target species by calling on the five functional modules in sequence. rENM() accepts a single four-letter bird banding code [55] as input. The code can represent any species that has been provisioned in the host environment’s data directory as described above.

**Table 7.** rENM package functions.

| Function | Description |
| --- | --- |
| rENM() | Execute the complete rENM pipeline for a target species |

In its current implementation, the rENM operates as a deterministic, unsupervised workflow controller: it executes each pipeline stage in a fixed sequence with no branching, conditional logic, or human intervention, making runs fully reproducible and providing a complete provenance record for each analysis through the run’s log file. Users who prefer finer control over individual stages, or who wish to run only a subset of the pipeline, can call the underlying module functions directly. The automated workflow uses default parameter settings across all package functions, and progress and timing for each stage are logged to a run-specific log file in the project directory (Fig 2).

### Framework usage

#### Interactive User Manual

The rENM Framework is distributed with an interactive User Manual implemented as a Quarto Markdown document [38]. Quarto is a scientific publishing system that combines executable code blocks and narrative text into a single, reproducible document [56]. Within the RStudio environment, the manual supports three modes of use. In the first, a user sets the four-letter banding code for a target species and steps through the document sequentially, reading the narrative description of each processing step and executing the accompanying code block before moving on. This mode is well suited to users who are learning the framework, as each section explains the purpose, options, and expected outputs of the corresponding workflow stage before any code is run. A user can also copy the user manual’s Quarto (.qmd) file, rename it, and use it as the starting point for a customized, species-specific analysis notebook fully edited by the user. Because the document combines function calls, parameter settings, and interpretive notes in a single file, it can serve alongside a run’s output directory as a complete and portable record of the analysis, one that can be shared with a publication as a companion to the reported results.

Finally, the entire workflow executes automatically when the user selects *Restart R* and *Run All Chunks* from the RStudio *Run* menu. This causes all code blocks to be processed in sequence without manual intervention. This batch execution mode produces the same outputs as the step-by-step mode and is equivalent in behavior to calling rENM() directly from the console.

Regardless of how the document is used, RStudio’s Quarto rendering engine can convert the Quarto files into multiple output formats from the *Render* drop-down menu, including .html, .pdf, and .docx. A Jupyter Notebook (.ipynb) version can also be rendered, producing a notebook that executes the same rENM Framework functions in an R kernel environment, making the workflow accessible to users who prefer the Jupyter interface without requiring any modification to the underlying R code.

### Model data

The rENM Framework is distributed with an example dataset that supports initial configuration, workflow familiarization, and testing. The dataset is available from the project’s Zenodo archive [57] and is downloaded and installed in the host environment’s `data` directory using a small set of shell commands provided in the User Manual (Fig 2). The example dataset includes occurrence records for three North American bird species: Brown-capped Rosy-Finch (*Leucosticte australis*; BCRF), Cassin’s Sparrow (*Peucaea cassinii*; CASP), and Greater Roadrunner (*Geococcyx californianus*; GRRO). These species were selected to represent various ecological contexts, range sizes, and occurrence record densities, thus allowing the user to explore how the framework behaves under different modeling conditions. To keep the example dataset lightweight and easy to distribute, we include only seven MERRA-2 microclimatic variables and seven MERRAclim-2 bioclimatic variables in the example dataset (see Schnase et al. [2] and S1 Appendix. MERRA-2 and MERRAclim-2 Variables for details). USGS GAP range polygons for each of the three species are also included in the example dataset. The example dataset is approximately 725 MB in size.

A more comprehensive dataset (∼75 GB), containing 30 MERRA-2 variables and the complete set of 19 MERRAclim-2 variables, is available separately for users wishing to extend beyond the example workflow and apply the framework to their own research questions (see the Data Availability Statement below for details). To smooth the representation of local environmental conditions, MERRA-2 and MERRAclim-2 variables in both the example and the comprehensive collections have been resampled from MERRA-2’s native spatial resolution of 1/2° latitude × 5/8° longitude to 5.0 arc-minutes (1/12°) resolution (∼7.6 km at latitude 35.0°N), as described in Schnase et al. [1]. The User Manual describes how users can extend the species occurrence records, environmental predictor collections, and range files to tailor the framework for their own work [38].

### Use of AI-Assisted Tools

The rENM Framework code base was developed with programming and documentation assistance from Claude Code [58] using the Claude Sonnet 4.6 model (Anthropic) [59]. The authors used Claude Sonnet 4.6 (Anthropic) [59] and GPT-5.5 (OpenAI) [60] as AI writing assistants to help with drafting, editing, and refining the manuscript text. All AI-assisted content was reviewed and edited by the authors, who independently verified the accuracy of the final text against source materials. The authors take full responsibility for the accuracy and integrity of the published work.

## Results

Because the rENM workflow incorporates stochastic procedures, individual results will vary to some degree among runs. In this section, we illustrate the general behavior of the framework and describe highlights from a representative run using Cassin’s Sparrow, the focal species used throughout framework development and the subject of previously published rENM studies [1,2,34–37]. The complete workflow was executed by calling `rENM("CASP")` on an Apple Mac Studio (M2 Ultra, 24-core, 128 GB RAM), producing outputs across all stages of the processing pipeline in 19.7 minutes of wall-clock time.

The complete output package comprised nearly 1,300 files totaling approximately 628.4 MB, including the run’s input occurrence records and environmental predictors, analysis products, logs, and performance diagnostics. Analysis products alone consisted of 181 files.

Together, these files constitute a complete provenance record of the run, sufficient to support independent verification and replication when executed within the rENM Framework’s R software environment. This example run is archived on Zenodo and can be downloaded in its entirety as a compressed archive file alongside runs for the two additional bird species for which example datasets are included in the current release of the framework [61]. The framework-generated final report for the example Cassin’s Sparrow run is provided as a supplementary appendix (S2 Appendix. rENM Framework example report) and can also be downloaded from the Zenodo archive along with reports for the other two example species [62].

### Performance

#### Data preparation and occurrence records

The eBird EBD for Cassin’s Sparrow contained 95,251 raw records, of which 88,747 had valid geographic coordinates [63]. After binning into nine five-year temporal intervals spanning 1980–2024, removing exact-coordinate duplicates, applying 1-km spatial thinning via `thin_occurrences2()`, and capping bins at 250 records, the final dataset comprised 2,206 occurrence records distributed across the nine bins. Record counts reflect the relative sparsity of early citizen-science data: the 1980–1984 bin retained only 206 records after thinning, while all subsequent bins reached the 250-record cap. This pattern of increasing record density through time is common across eBird-supported taxa and may have implications for ensemble model stability in earlier time steps that users should be aware of. The USGS GAP range polygon for CASP was used to define the spatial extent (−115° to −94°W, 26° to 43°N), encompassing a modeled area of approximately 3.62 million km² with a range footprint of 1.45 million km².

### Variable screening and time-series construction

The convergence-based variable screener (`screen_by_convergence2()`) was applied independently to each of the nine five-year bins against the complete complement of 49 candidate MERRA-2 and MERRAclime-2 variables, resolving a top-*n* = 7 variable set for each bin. Convergence was rapid and consistent, ranging from 735 predictor-pair evaluations (1990 and 2010 bins) to 3,675 predictor-pair evaluations (1980 bin), with elapsed screening times per bin ranging from under one minute to approximately 2.25 minutes. The fastest convergence occurred in bins with higher occurrence record density, consistent with evidence that smaller training samples reduce the reliability of variable importance estimation [64,65]. The top 7 variables per bin were then staged for ensemble modeling.

The full ensemble time series was constructed by create_timeseries(), which executed nine five-year bin models in parallel, each fitting five algorithms (maxnet, RF, BRT, GLM, and MARS) with three replicates and 2,500 background points. Model performance was consistently high across the nine time bins (1980–2020). All 135 individual model runs (five methods × nine bins × three replicates) achieved area under the receiver operating characteristic curve (AUC) > 0.70 on withheld test data, with a grand mean AUC of 0.845 (range: 0.72–0.92). Following established thresholds, AUC values in this range indicate good to excellent discriminative ability [49,66], with ensemble mean AUC per bin ranging from 0.822 to 0.874. True Skill Statistic (TSS) values (ensemble mean ∼0.58) corroborated these assessments [67]. The operation completed 135 total model fits in approximately 2.5 minutes. Each bin produced continuous suitability and binary range rasters as primary outputs, which were subsequently consumed by the analysis module.

### Analytical outputs

#### Output products

For each run, `rENM.analysis` produced five classes of analytical outputs, illustrated here for the Cassin’s Sparrow example with sample pages from the framework-generated analysis report (Fig. 3). The report opens with an AI-generated synthesis of the run’s quantitative results, followed by a reference list and AI disclaimer. A nine-panel suitability time series and corresponding range map panel show the evolving spatial structure of modeled climatic suitability across the study period. A suitability trend map shows the direction and magnitude of long-term change across the modeled extent, with movement of the suitability-weighted centroid overlaid to show the net directional shift; a companion map identifies regions of accelerating or decelerating suitability change. State-level trend and hotspot maps summarize the geographic distribution of suitability change and areas of accelerating decline within the species’ USGS GAP range. Centroid displacement plots show the Bayesian-estimated longitudinal and latitudinal shift in the weighted centroid for suitability over the 45-year study period.

**Fig. 3.**
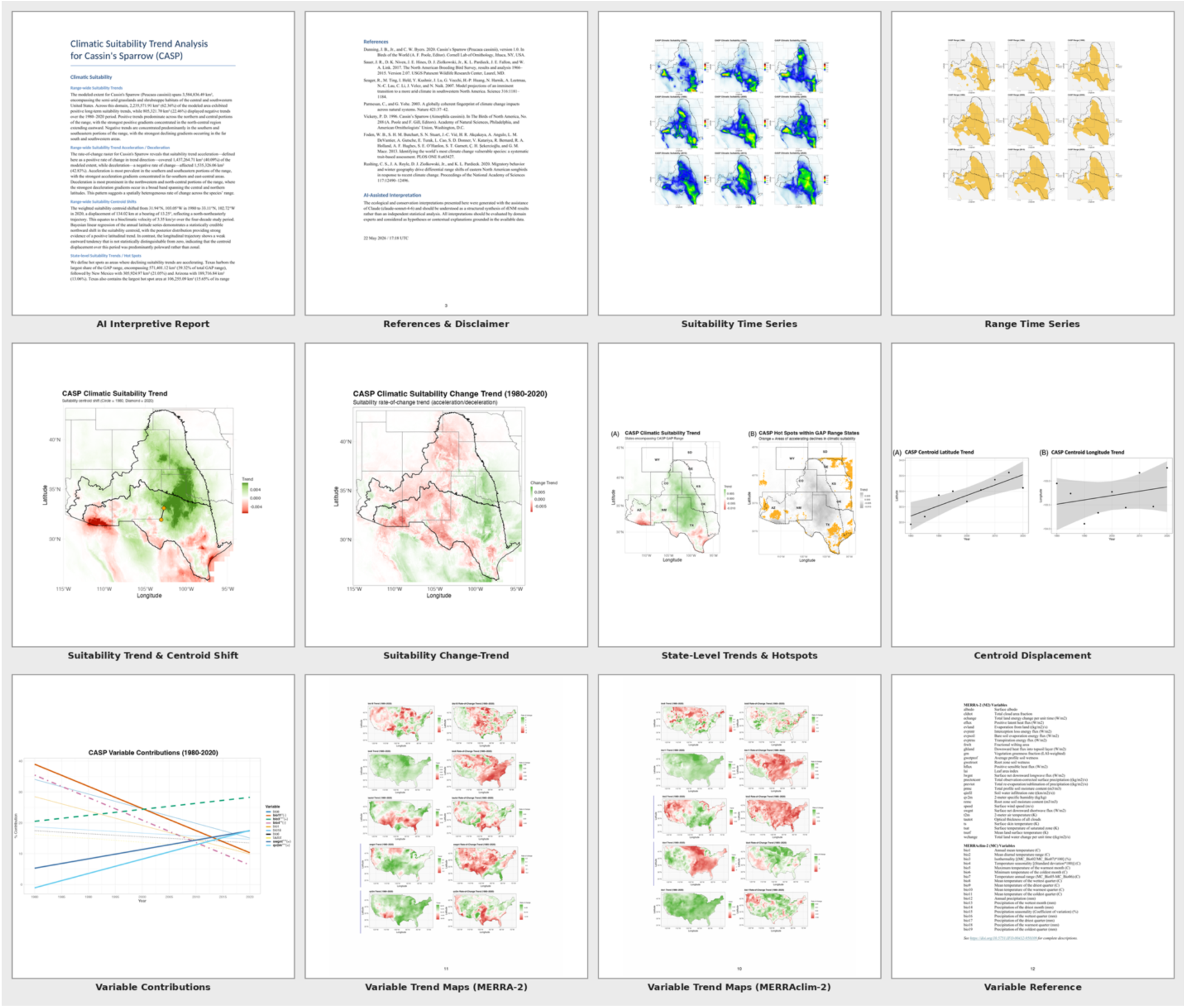
Representative pages from an rENM Framework analysis report. Image shows the scope and diversity of outputs generated by a single pipeline run for Cassin’s Sparrow. Row 1: AI-generated interpretive narrative; automatically generated references and AI disclaimer; ensemble climatic suitability time series; range map time series. Row 2: Theil-Sen suitability trend map with 45-year centroid shift; suitability acceleration/deceleration map; state-level trend and hotspot maps; 45-year longitude/latitude centroid displacement plots. Row 3: variable contribution trends; variable trend maps for the top 10 MERRA-2 predictors in the overall time series; variable trend maps for the top 10 MERRAclim-2 predictors in the overall time series; variable reference guide.

Variable contribution trend plots show how the relative importance of the top environmental predictors changes across the nine temporal bins, with accompanying Bayesian regression statistics. Variable trend maps show the spatial distribution of change in the top MERRA-2 and MERRAclim-2 predictors themselves over the past 45 years, thereby allowing quick visualization of the dynamic nature of these model influences. Finally, a variable reference guide describes the suite of variables used in the run. Together these products provide an analytical portrait of a species’ multi-decadal response to changing climatic conditions.

### AI-assisted interpretation

The rENM Framework currently supports two experimental options for obtaining AI-assisted analytical interpretations. For comparison, we obtained results from both options in this example. As a final stage of the processing pipeline, `rENM.ai` automatically submitted a set of quantitative outputs from the Cassin’s Sparrow run to Claude (claude-sonnet-4-6) and ChatGPT (gpt-5.1). ChatGPT completed its response in 132 seconds at a cost of <$>0.10\semicolon \; Claude required 450 seconds at a cost of <$>2.29. Both reports covered the same analytical domains: range-wide suitability trends, acceleration and deceleration, centroid displacement, and state-level hotspots. Each also appended an ecological and conservation interpretation section with an automatically generated reference list and a standardized disclaimer that interpretations represent data-grounded hypotheses rather than validated scientific conclusions and should be evaluated by domain experts. Both correctly identified the major quantitative results of the run and connected the hotspot geography to known conservation concerns for the species [2,68].

The reports differed in emphasis: Claude’s was more quantitatively precise and offered more specific ecological interpretation of individual variable trends, while ChatGPT’s was more concise and placed greater emphasis on actionable conservation management recommendations.

These differences underscore the importance of critical expert review before AI-generated interpretations inform conservation decisions. Despite being generated through independent API calls with no shared context, the two reports provided broadly similar high-level interpretations. This convergence suggests that the inputs produced by the rENM pipeline constrain the narrative space sufficiently that different AI models arrive at comparable syntheses, an observation that further suggests AI-assisted interpretation might provide a reproducibility check on complex, automated workflows such as that seen in the rENM Framework.

## Discussion

Our results do not provide a formal validation of rENM or any of the individual components of the rENM Framework. What they do represent is a growing body of evidence that the approach generates analytical products that complement established methodologies in useful ways and that open lines of ecological inquiry that conventional ENM approaches cannot easily address. In the sections that follow, we describe the areas where we see the most promising contributions and practical advances coming out of this work and the opportunities for refinement we think the community could help address, including our plans for future development.

### Framework contributions

#### Applications to species ecology

Successive analyses over the past several years tell a coherent and increasingly detailed story about Cassin’s Sparrow’s climatic ecology that would have been inaccessible without the time-series approach described here. Our earliest use of rENM demonstrated that modeled estimates of climatic suitability for the species vary substantially depending on the temporal frame, spatial extent, and environmental drivers considered and that this variability mirrors the inconsistent portrait of the species’ conservation status that has persisted in the literature for decades [1]. We then used rENM to examine the intra-annual dynamics of suitability across Cassin’s Sparrow’s breeding season. This work revealed an east-to-west progression in suitable conditions that aligns with longstanding hypotheses about the species’ opportunistic breeding movements [2]. The study also documented a decline in the distance of these movements that suggests the species may be adapting to regional environmental conditions rather than making the range-wide movements long thought to define its breeding strategy, a finding we regard as among the most ecologically significant to emerge from this body of work.

Analysis of variable contributions has also helped characterize the environmental structure of Cassin’s Sparrow’s niche and how that structure has evolved over time. Across our time series models, drying-related energy flux variables have generally gained explanatory weight while moisture availability terms have declined [1,2]. This result is consistent with regional warming and aridification trends in the southwestern United States [69,70] and with the broader recognition that species-environment relationships can shift in complex ways across time and space [20,21].

We regard this consistency across successive studies, each employing different data sources, predictor collections, and temporal resolutions, as a demonstration that rENM is detecting real ecological patterns rather than artifacts of any particular methodological configuration. It also gives us confidence that the techniques described here have broader applicability.

### Applications to conservation practitioners

State Wildlife Action Plans (SWAPs) provide the primary framework through which state fish and wildlife agencies identify and prioritize conservation activities for Species of Greatest Conservation Need (SGCN) [71,72]. The picture that emerges from the SWAPs of the four states that encompass the core of Cassin’s Sparrow’s US breeding range suggests that conservation applications may represent rENM’s most consequential future use.

New Mexico lists Cassin’s Sparrow as a Category D (Data Needs) SGCN carrying climate change vulnerability, decline, and vulnerability flags across multiple ecoregions [73]. Arizona lists it as a Tier 3 SGCN, the tier reserved for species whose status cannot be adequately assessed due to a lack of information [74]. Colorado does not list the species as an SGCN [75]. Texas, which encompasses a substantial portion of the species’ US breeding range, does not list it either [76]. This four-state pattern of inconsistent SGCN status, variable data gaps, and no shared analytical framework for assessing species’ status across state boundaries is not unusual. States differ in administrative procedures, biodiversity data platforms, taxonomic standards, SGCN selection criteria, and capacity to implement conservation, producing SGCN lists and status designations that vary substantially across states, particularly for wide-ranging species with distributions spanning multiple states [77,78]. The Cassin’s Sparrow case is an example of a type of challenge that recurs across the national SWAP effort.

The dominant tool currently available for climate vulnerability assessment in the SWAP context is the NatureServe Climate Change Vulnerability Index (CCVI), which generates a categorical score by integrating a species’ exposure to projected climate change with factors related to sensitivity and adaptive capacity [79]. New Mexico applied CCVI Version 4.0 to 295 vertebrate SGCN in its 2025 SWAP revision [73]. The index scores each species under two of the Intergovernmental Panel on Climate Change (IPCC) Representative Concentration Pathway (RCP) greenhouse gas concentration scenarios [80] (RCP 4.5, representing moderate mitigation of greenhouse gas emissions, and RCP 8.5, representing a high-emissions trajectory) to bracket the range of plausible climate futures. Cassin’s Sparrow scored as Less Vulnerable under both and was not alone: at least 71% of bird SGCN in that same analysis scored as Less Vulnerable. This pattern is not specific to New Mexico. Migratory birds tend to score well in the CCVI’s adaptive capacity component, because high dispersal ability is treated as a buffer against climate exposure; more fundamentally, the index assesses vulnerability only during the breeding season, which means threats faced on migration and wintering grounds are not captured [81] . This is not a criticism of the CCVI; it performs as designed, and it is a well-established tool for rapid, cost-effective screening. But a "Less Vulnerable" score for a species simultaneously flagged for decline, climate change vulnerability, and data deficiency in two states and not considered to be SGCN in two others illustrates the gap the rENM Framework is positioned to fill.

The CCVI produces categorical summaries. While it can demonstrate how a species’ vulnerability differs from state to state, it cannot resolve where within a state or a species’ range conditions are deteriorating, or how rapidly. The rENM Framework operates at this finer resolution. Among its outputs, the hotspot analysis, which identifies areas of accelerating suitability decline within a species’ range, may prove to be the most directly actionable product for conservation practitioners. No static ENM produces this output, and few tools in the current SWAP toolkit generate a reproducible basis for prioritizing species-specific monitoring intensity and management attention at those sub-state spatial scales. The hotspot analysis has not yet been validated against independent field data and should be treated as a hypothesis-generating product pending that evaluation. However, we believe its potential utility for targeting survey effort and informing within-range management decisions is significant.

The second contribution the framework makes to conservation practice is one of scalability. State wildlife agencies face persistent resource constraints that limit how many species they can study in depth; for many SGCN, and particularly those assigned to data-deficiency categories such as New Mexico’s Category D or Arizona’s Tier 3, the lack of biological information is itself the central conservation problem. The rENM Framework’s automated, unsupervised pipeline changes what is practically achievable. Because a complete analysis can be executed for any provisioned species in a matter of minutes, systematic application across large numbers of SGCN becomes tractable for the first time. This reduces cost, expands species coverage, and makes cross-species comparison straightforward. Scalability also makes guild-level analysis possible. When suitability trajectories can be generated simultaneously for all grassland sparrows, or all arid-land obligates, or all species sharing a migratory strategy, patterns invisible at the single-species level may emerge, adding a dimension to conservation planning that state and regional SWAP processes currently cannot access, such as identifying state-level or regional areas where hotspots of suitability decline overlap across multiple species and therefore where on-the-ground restoration efforts may be needed.

### Opportunities for refinement

#### Data provisioning

There are two areas where enhanced data provisioning would increase the framework’s analytical power. First, the current framework draws its environmental predictors entirely from MERRA-2 and its derived MERRAclim-2 bioclimatic variables. This is an important data source: MERRA-2’s temporal depth and internal consistency are what make the 45-year time series possible. But climate variables alone cannot fully characterize a species’ habitat, and several classes of data could meaningfully expand the framework’s explanatory reach.

The most useful near-term addition is vegetation. For example, Cassin’s Sparrow is a grassland-shrubland obligate whose distribution appears to track fine-scale changes in vegetation structure. Broad-scale climatic variability, fine-scale vegetation, and topographic heterogeneity drive habitat suitability trends in grassland and shrubland systems [82,83]. Incorporating a vegetation time series that is temporally co-registered with MERRA-2 climate variables would move the framework toward modeling habitat suitability in a more complete sense. Soil properties, such as texture and organic matter content, influence vegetation composition and structure and are also important in arid and semi-arid systems. Soil datasets accompanied by hydrological variables could capture precipitation-driven habitat dynamics that matter for species like Cassin’s Sparrow that track moisture availability closely. Land cover and land use change could provide a structural layer that distinguishes developed, agricultural, and natural cover at resolutions appropriate for range-scale analyses.

A second important data provisioning enhancement involves occurrence records. The framework currently ingests eBird EBD files and is therefore effectively limited to analyses of the environmental associations of birds. Extending the occurrence data to accommodate other sources would expand the framework’s taxonomic reach and facilitate the guild-level and multi-taxa analyses described above. The most immediate targets are well-established platforms with large, openly accessible, research-grade observational archives. The Global Biodiversity Information Facility (GBIF) [84] aggregates occurrence records across all taxa from hundreds of contributing institutions and citizen-science networks, making it the most comprehensive single access point for non-avian species. iNaturalist [85] provides particularly dense coverage for invertebrates, reptiles, amphibians, and plants. For taxa with dedicated monitoring communities, more specialized sources may be possible, such as eButterfly [86] and the Botanical Information and Ecology Network (BIEN) database [87] for butterflies and New World plants, respectively. Each of these additions would require evaluation for temporal availability and the capacity for co-registration with existing environmental data collections, as well as functional refinements to the rENM.data module before data from these other sources can be analyzed through the framework.

### Modeling and analysis

The two most consequential modeling and analysis enhancements on our agenda are forward projection and higher-resolution temporal analysis. Forward projection from a retrospective environmental suitability time series baseline is, to our knowledge, new ground for the field of ENM. Conventional ENM projections calibrate models against a single present-day snapshot and propagate those relationships into future climate scenarios. The limitations to this approach are well established: when species-environment relationships are demonstrably non-stationary, a single baseline may misrepresent the trajectory that future projections should extend [28–30]. A 45-year retrospective time series provides a record of directional change and a fundamentally different departure point for forward projection. The question is how to use that record.

We see two candidate approaches, both currently speculative. The first would use the suitability trend and acceleration/deceleration maps produced by the framework’s rENM.analysis step as the starting states for projection rather than a lone suitability map. These trend maps encode not just suitability at a point in time but how the direction and rate of suitability has changed over the past 45 years. Projecting forward from a trend rather than a single state is conceptually closer to extrapolating a trajectory than to transferring a niche, and it may be especially useful for species undergoing niche reorganization. The second approach would use the temporal trajectory of changes in variable importance across the modeled time series to identify which projected climate variables are most consistent with the drivers of historical suitability change. Rather than projecting a static niche model into a future climate state, this approach would use the trajectory of changes in environmental relationships documented in the retrospective record to identify which projected future conditions are most ecologically relevant to the species. The framework already generates the variable contribution trend record this approach would draw on; what remains to be developed is the methodology for matching that record against global climate model projections, evaluating model performance, and building the codebase to support variable selection and projected model results.

The second development priority is higher temporal resolution time series analyses. The current framework operates at five-year intervals, which captures long-term trends effectively but cannot resolve the intra-annual dynamics that matter for many species. Schnase et al. [2] demonstrate the value of a finer-grained, seasonal approach. That work constructed a biweekly time series for Cassin’s Sparrow’s breeding season that revealed seasonal suitability dynamics over the past 40 years that are invisible at the five-year resolution. The research codebase supporting that work is available for integration into a future version release of the rENM Framework. That integration will require substantial functional modifications across the entire framework: the temporal binning logic, predictor staging procedures, model execution pipeline, and output summarization tools all assume five-year intervals and will need to be generalized. The payoff will be access to a class of ecological questions about seasonal range dynamics that the current framework cannot address.

### AI interpretation and agentic control

The application of large language models (LLMs) to ecological research is an active and rapidly developing area. Published work has explored LLMs across several tasks relevant to biodiversity science: extracting taxonomic and geographic information from ecological literature [88], using language-driven taxonomic embeddings to improve SDM performance [89], and evaluating the factual ecological knowledge that off-the-shelf LLMs carry, including species presence prediction, range mapping, and threat classification [90]. The closest analogue to what rENM.ai does is the work of Gallois et al. [91], who present a roadmap for building bespoke quantitative LLMs as virtual ecological data assistants, designed to transform structured observational datasets into interpretive outputs and actionable conservation insights. What distinguishes rENM.ai from all of these efforts is not the use of LLMs per se but the context in which they operate. The model does not retrieve or classify information from text, nor does it answer natural language queries about species ranges. It receives a curated, quantitative data bundle assembled automatically from a completed analytical run and returns a structured interpretive narrative as output.

The current implementation operates at the workflow level: a single LLM prompt submitted after a full, multi-step analysis is completed summarizes the run’s outputs across all of the framework’s modules. This is a useful starting point, but a more powerful approach would deploy LLMs at the step level, submitting prompts as each analytical module completes. This would allow the LLM to reason about, and perhaps correct, intermediate results before the full analysis is assembled. Step-level prompting could support more nuanced interpretation and could enable a form of iterative, adaptive refinement of the overall analysis that the current end-of- pipeline LLM synthesis cannot produce.

The second potential AI frontier is the rENM Framework’s capacity for autonomous agentic control. Autonomous agentic control refers to the capacity of AI-driven systems to independently orchestrate complex, multi-step scientific workflows, such as selecting tasks, allocating resources, and adapting to intermediate results, with minimal or no human intervention at each stage [92]. In its current form, rENM processing executes pipeline stages in a fixed sequence with no branching, no feedback, and no adaptive behavior. This determinism makes runs fully reproducible and provides a complete provenance record, important elements of scientific documentation. It also limits what the system can do on its own.

A next step in the evolution of the rENM methodology would involve adding an AI-driven agentic controller that autonomously supervises the behavior of the entire pipeline, essentially moving the framework from a static system toward adaptive and optimizing behavior [83]. A controller such as this would hierarchically coordinate the activities of the lower-level, agentically controlled modules: selecting which species to analyze based on conservation priority criteria, acquiring inputs, triggering analyses when new occurrence data or updated environmental predictor collections arrive in monitored data sources, comparing outputs across species to identify emergent patterns, and routing results to appropriate downstream consumers such as SWAP databases or agency monitoring dashboards. Autonomous monitoring of SGCN across a state’s full species list, periodic re-analysis triggered by updated eBird data, and systematic detection of accelerating suitability declines without human initiation are all plausible applications of this kind of system. The implications for conservation practice are substantial: a state wildlife agency equipped with an autonomous rENM monitoring pipeline would have analytical capacity that no current manual workflow can match. We regard the integration of AI into the science and application of rENM as the most consequential direction for future development, but one whose transformational potential must be weighed against the environmental costs of the infrastructure it depends on.

This environmental cost is not incidental. Data centers and AI infrastructure carry substantial and growing energy and water demands [93,94], and these costs are not evenly distributed [95]. The same arid landscapes that are home to Cassin’s Sparrow and many other SGCN and that are particularly vulnerable to climatic change, are, in many cases, the landscapes where new AI infrastructure is being aggressively developed and where its water and energy demands will be most severely felt. Data center expansion across the arid Southwest, including New Mexico, Colorado, Arizona, and Texas, is colliding directly with water scarcity in a region already defined by drought and an over-allocated Colorado River system [96,97]. Realizing AI’s potential for informing conservation science will require confronting this tension directly rather than treating computational capability as costless [98]. We do not resolve that tension here, but we believe it deserves explicit acknowledgment in any discussion of AI’s future role in ecological modeling and conservation practice. We cannot afford to continually degrade the environments we are trying to help by using the modeling tools described here.

## Conclusions

The rENM Framework is a modular, open-source suite of R packages that reconstructs and analyzes long-term ecological niche dynamics by combining historical bird occurrence records with a 45-year time series of NASA MERRA-2 environmental data. The framework automates a complete retrospective ecological niche modeling workflow, from data preparation and ensemble time-series construction through trend analysis, AI-assisted interpretation, and report generation and executes it for any species for which the user has provided occurrence and environmental predictor data with a single function call.

The framework addresses a well-recognized shortcoming in conventional ENM practice: the tendency to represent species-environment relationships as static, time-averaged summaries that cannot detect nonstationarity or reveal how climatic suitability has changed across decades. By treating the evolving suitability surface as an object of study rather than a fixed baseline, rENM produces analytical products that complement established methods and open lines of ecological inquiry that static approaches cannot easily address. Illustrated here with Cassin’s Sparrow, a grassland species of conservation concern in the arid southwestern United States, those products include suitability time series, trend and acceleration maps, centroid displacement estimates, bioclimatic velocity metrics, variable contribution trajectories, and hotspot analyses identifying areas of accelerating suitability decline.

The framework’s automated, unsupervised pipeline makes systematic application across large numbers of species tractable for the first time, with direct implications for state wildlife agencies managing Species of Greatest Conservation Need under resource constraints. The experimental AI-assisted interpretation module demonstrates that large language models can function as embedded workflow components rather than standalone tools, generating structured ecological narratives from curated quantitative outputs. Both the analytical products and the AI integration remain experimental; validation against independent field data and broader taxonomic testing will be essential next steps. The most important directions for future development are forward projection from a retrospective baseline, higher temporal resolution, and the extension of the current automated pipeline toward autonomous agentic control, a progression that could give conservation agencies analytical capacity that no current manual workflow can match. The framework is openly available on GitHub and archived on Zenodo, with example datasets and an interactive user manual, and is ready for application, evaluation, and further community development.

## Supporting Information

**S1 Appendix. MERRA-2 and MERRAclim-2 variables**. Overview of the MERRA-2 variables used in the rENM Framework Version 0.1.0 release.

**S2 Appendix. rENM Framework report**. Example rENM analysis report for Cassin’s Sparrow.

## Data Availability Statement

The rENM Framework source code is openly available as a suite of R packages on GitHub [39] and archived on Zenodo [40,99] along with a User Manual [38] and individual package archives for rENM [100], rENM.core [101], rENM.data [102], rENM.model [103], rENM.analysis [104], rENM.ai [105], and rENM.reports [106]. The example input dataset required to reproduce the analysis described in this paper is archived on Zenodo [57]. Complete output runs and reports for the three example species are archived on Zenodo [61,62], along with the extended MERRA-2 and MERRAclim-2 data collection [107]. Information on extending the framework, including how to download data for other species, how to acquire USGS GAP range files, and adding new variables types is provided in the User Manual [38].

## Author contributions

JLS: Conceptualization, Methodology, Investigation, Software, Formal analysis, Writing – original draft. MLC, PMM, VAS: Conceptualization, Methodology, Formal analysis, Writing – review & editing.

### S1 Appendix. MERRA-2 and MERRAclim-2 variables

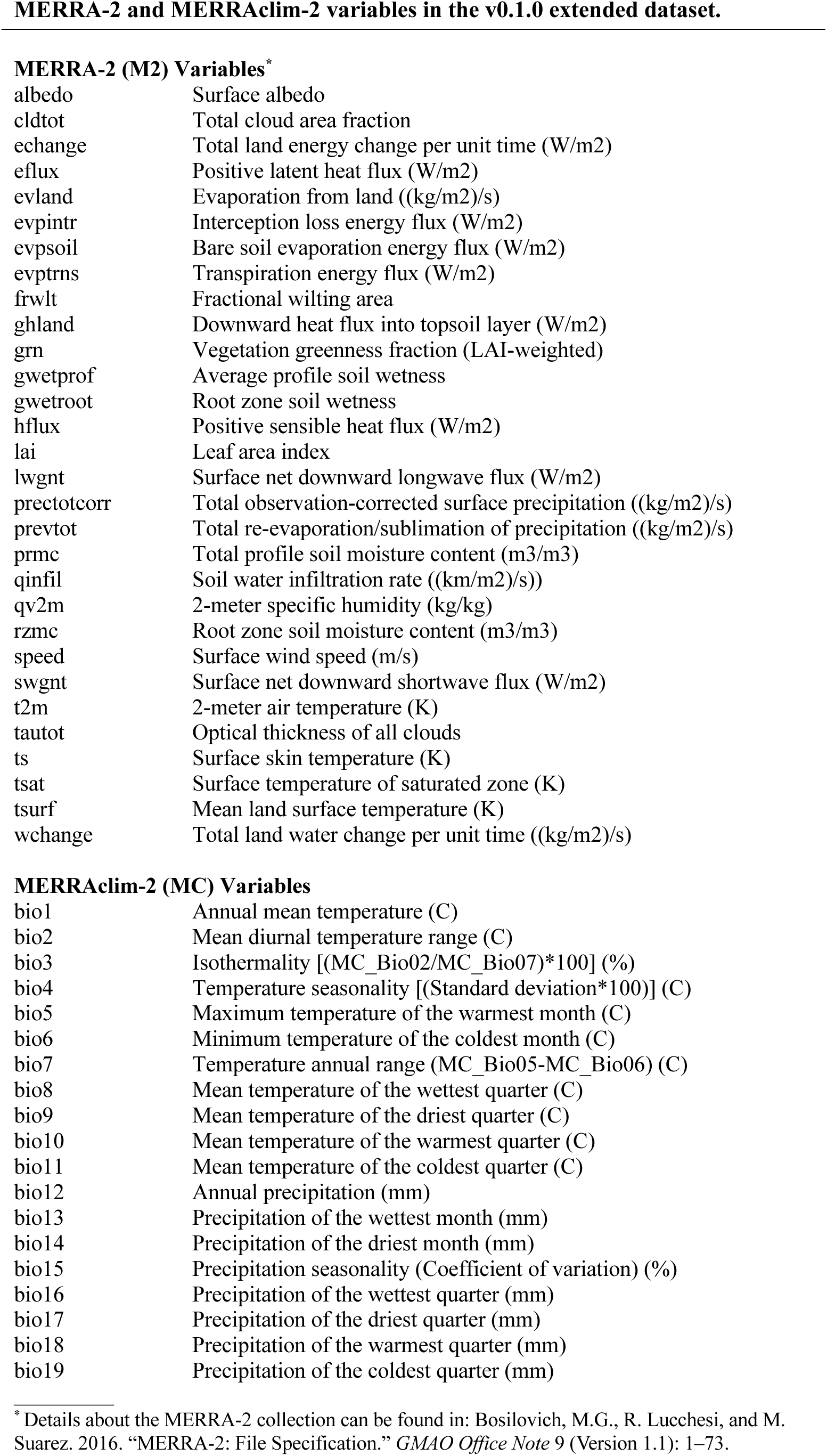

MERRA-2 (M2) Variables\*
|  |  |
| --- | --- |
| albedo | Surface albedo |
| cldtot | Total cloud area fraction |
| echange | Total land energy change per unit time (W/m2) |
| eflux | Positive latent heat flux (W/m2) |
| evland | Evaporation from land ((kg/m2)/s) |
| evpintr | Interception loss energy flux (W/m2) |
| evpsoil | Bare soil evaporation energy flux (W/m2) |
| evptrns | Transpiration energy flux (W/m2) |
| frwlt | Fractional wilting area |
| ghland | Downward heat flux into topsoil layer (W/m2) |
| grn | Vegetation greenness fraction (LAI-weighted) |
| gwetprof | Average profile soil wetness |
| gwetroot | Root zone soil wetness |
| hflux | Positive sensible heat flux (W/m2) |
| lai | Leaf area index |
| lwgnt | Surface net downward longwave flux (W/m2) |
| prectotcorr | Total observation-corrected surface precipitation ((kg/m2)/s) |
| prevtot | Total re-evaporation/sublimation of precipitation ((kg/m2)/s) |
| prmc | Total profile soil moisture content (m3/m3) |
| qinfil | Soil water infiltration rate ((km/m2)/s)) |
| qv2m | 2-meter specific humidity (kg/kg) |
| rzmc | Root zone soil moisture content (m3/m3) |
| speed | Surface wind speed (m/s) |
| swgnt | Surface net downward shortwave flux (W/m2) |
| t2m | 2-meter air temperature (K) |
| tautot | Optical thickness of all clouds |
| ts | Surface skin temperature (K) |
| tsat | Surface temperature of saturated zone (K) |
| tsurf | Mean land surface temperature (K) |
| wchange | Total land water change per unit time ((kg/m2)/s) |
| MERRAclim-2 (MC) Variables |  |
| bio1 | Annual mean temperature (C) |
| bio2 | Mean diurnal temperature range (C) |
| bio3 | Isothermality [(MC_Bio02/MC_Bio07)*100] (%) |
| bio4 | Temperature seasonality [(Standard deviation*100)] (C) |
| bio5 | Maximum temperature of the warmest month (C) |
| bio6 | Minimum temperature of the coldest month (C) |
| bio7 | Temperature annual range (MC_Bio05-MC_Bio06) (C) |
| bio8 | Mean temperature of the wettest quarter (C) |
| bio9 | Mean temperature of the driest quarter (C) |
| bio10 | Mean temperature of the warmest quarter (C) |
| bio11 | Mean temperature of the coldest quarter (C) |
| bio12 | Annual precipitation (mm) |
| bio13 | Precipitation of the wettest month (mm) |
| bio14 | Precipitation of the driest month (mm) |
| bio15 | Precipitation seasonality (Coefficient of variation) (%) |
| bio16 | Precipitation of the wettest quarter (mm) |
| bio17 | Precipitation of the driest quarter (mm) |
| bio18 | Precipitation of the warmest quarter (mm) |
| bio19 | Precipitation of the coldest quarter (mm) |

\*Details about the MERRA-2 collection can be found in: Bosilovich, M.G., R. Lucchesi, and M. Suarez. 2016. "MERRA-2: File Specification." *GMAO Office Note 9* (Version 1.1): 1–73.

### S2 Appendix. rENM Framework report

#### Climatic Suitability Trend Analysis for Cassin’s Sparrow (CASP) 1980–2024

##### CLIMATIC SUITABILITY

###### Range-wide Suitability Trends

The Theil–Sen trend surface indicates substantial spatial structure in long-term climatic suitability for Cassin’s Sparrow across a modeled extent of 1,838,111.52 km². Within this area, 1,207,694.06 km² (65.70%) show positive suitability trends, whereas 448,960.92 km² (24.43%) exhibit negative trends. The modeled extent spans the south-central United States, from the southern Great Plains northward into the central High Plains, encompassing portions of the desert Southwest and adjacent prairie regions. Positive suitability trends are concentrated across the central and eastern portion of the modeled range, especially through the central Great Plains and portions of the southern High Plains, whereas negative trends are more common toward the western and southwestern margins of the range.

#### Range-wide Suitability Trend Acceleration / Deceleration

The rate-of-change surface reveals pronounced contrasts between areas where suitability trends are accelerating and areas where they are decelerating. Acceleration, defined as an increasing rate of change in climatic suitability over time, covers 726,906.25 km² (39.55%) of the modeled extent, whereas deceleration, defined as a decreasing rate of change, occupies 921,467.98 km² (50.13%). Areas where suitability trends are accelerating are most extensive through the central and eastern sectors of the range, particularly across the southern and central Great Plains. Deceleration is more localized, occurring mainly along the western and southwestern edges of the modeled extent and in scattered patches toward the northern margin.

#### Range-wide Suitability Centroid Shihs

The bioclimatic velocity analysis indicates that the weighted climatic suitability centroid shifted from approximately 32.20°N, 102.76°W in 1980 to 32.95°N, 102.73°W in 2020, a displacement of 83.32 km along a 1.68° northward trajectory. This corresponds to an average bioclimatic velocity of 2.08 km/yr over the study period. Bayesian trend analysis of the annual centroids shows a statistically clear directional change in latitude and a statistically clear directional change in longitude across 1980–2020, indicating systematic movement of the suitability centroid in both the north–south and east–west dimensions without detailing specific effect sizes.

#### State-level Suitability Trends / Hot Spots

We define hot spots as areas where declining suitability trends are accelerating. Within the USGS GAP range, states differ markedly in their proportional contributions to the modeled distribution. TX contains 571,401.12 km² of GAP range, representing 39.32% of the total modeled range. NM contains 305,924.97 km² of GAP range, representing 21.05% of the total modeled range. AZ contains 189,716.84 km² of GAP range, representing 13.06% of the total modeled range. Hot spot coverage also varies among states. TX contains 58,876.75 km² of hot spots (11.93% of the state’s modeled range). AZ contains 12,961.04 km² of hot spots (12.52% of the state’s modeled range).

#### Ecological / Conservation Interpretation

For Cassin’s Sparrow, the predominance of positive suitability trends and substantial accelerating areas across the central and eastern portion of the modeled range suggest that climatic conditions in much of the southern and central Great Plains have recently become more compatible with the species’ realized niche. Northward centroid displacement indicates progressive alignment of suitable climates with higher latitudes, consistent with documented climate-driven shifts in grassland bird distributions. Concentrations of hot spots in Texas and Arizona highlight regions where rapid climatic deterioration may outpace the species’ capacity to track favorable conditions through dispersal or demographic compensation. These areas likely warrant heightened monitoring of population trends, habitat structure, and land-use change, given the known sensitivity of arid and semi-arid grasslands to altered precipitation regimes, increasing temperatures, and woody encroachment, and the recognized dependence of Cassin’s Sparrow on open, shrub–grass mosaics within these systems.

### ENVIRONMENTAL STRUCTURE

#### Variable Profile

Bayesian trend analyses of the top contributing environmental predictors show that several variables exhibit statistically important directional changes across the 1980–2020 period. Variables with non-overlapping regions of practical equivalence for their slope parameters include speed, bio4, bio10, bio11, bio8, lai, bio17, bio15, bio14, and qv2m, indicating consistent temporal trajectories in their modeled contributions. Collectively, these predictors describe gradients in temperature, precipitation, atmospheric moisture, vegetation structure, and surface wind speed across the southern Great Plains and adjacent arid landscapes. Over the study interval, the regression slopes indicate systematic shifts in these gradients, with some variables showing sustained increases and others exhibiting declines in their relative importance through time, producing a temporally evolving climatic and structural backdrop for suitability patterns.

#### Ecological / Conservation Interpretation

The combination of changing temperature regimes, precipitation seasonality, atmospheric moisture, and vegetation structure implied by the variable trends is consistent with widespread climatic reorganization across the southern Great Plains and Southwest during recent decades. Increases in the influence of warm-season temperature metrics and water-balance variables align with observations of intensified heat and more variable precipitation in North American drylands, while changing leaf area and wind speed patterns reflect interacting land-use and climatic drivers. For Cassin’s Sparrow, whose occurrence is closely associated with semi-arid shrub–grass mosaics, these structural shifts likely modify both the configuration and temporal stability of suitable habitat. Conservation strategies that integrate projected trajectories of key climatic and vegetation variables with ongoing grassland management and restoration efforts will therefore be important for sustaining populations under continuing environmental change.

## AI-ASSISTED INTERPRETATION

The ecological and conservation interpretations presented here were generated with the assistance of ChatGPT (GPT 5.1) and should be understood as a structured synthesis of rENM results rather than an independent statistical analysis. All interpretations should be evaluated by domain experts and considered as hypotheses or contextual explanations grounded in the available data.

## INCLUDED FIGURES

Page 5 - Climatic Suitability Time Series

Page 6 - Range Time Series

Page 7 - Climatic Suitability Trends

Page 8 - State-Level Suitability Trend and HotSpot Summary

Page 9 - Climatic Suitability Trend with Centroid Shift

Page 10 - Variable Contribution Trends

Page 11 - Predictor Variable Trends

Page 14 - rENM Framework MERRA Variables

**For additional information about the rENM Framework, see:**

Schnase, J. L., M. L. Carroll, P. M. Montesano, and V. A. Seamster. 2026. The rENM Framework: Toward a Modular System for Reconstructing and Analyzing Long-Term Ecological Niche Dynamics. bioRxiv preprint, in preparation.

(rENM Framework Version 0.1.0 - 30 July 2026 - 15:10 UTC)

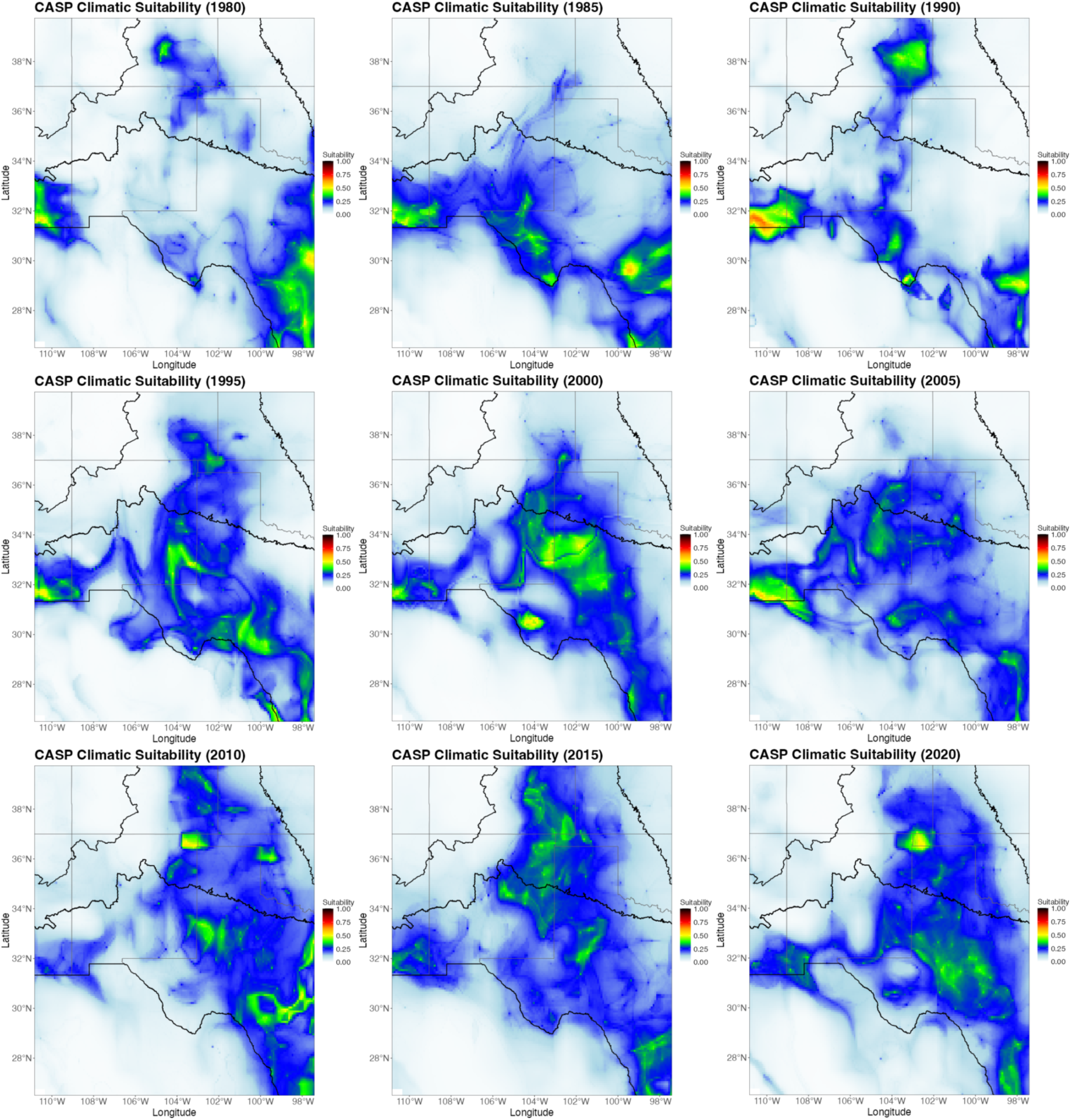

CLIMATIC SUITABILITY TIME SERIES. Each of the nine panels shows the ensemble-modeled climatic suitability surface for one temporal bin across the study period, arranged chronologically from left to right and top to bottom. Higher suitability values appear in warmer colors (yellow, green); lower values appear in cooler colors (blue, white). The black outline delineates the USGS GAP range boundary for the species.

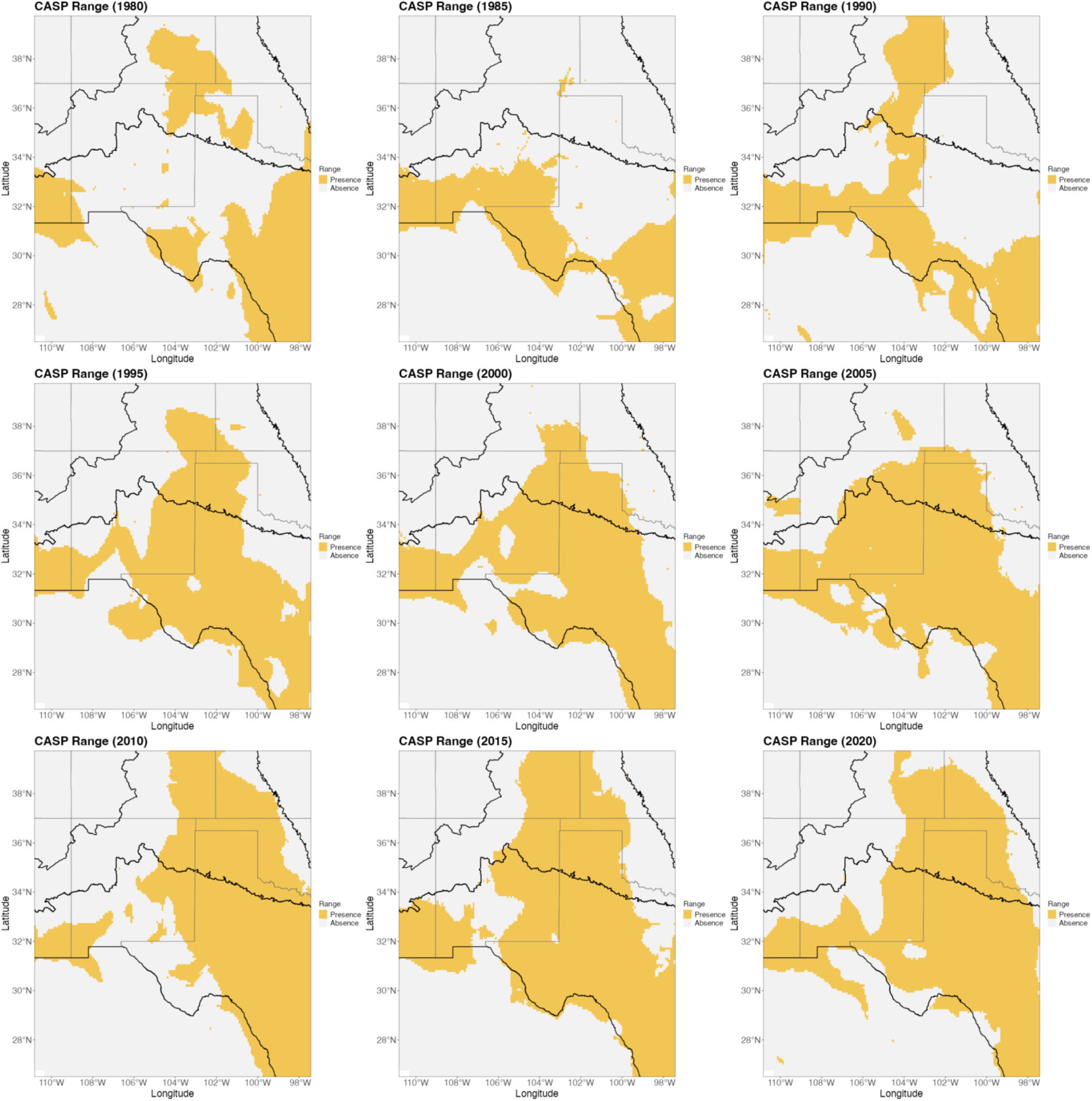

RANGE TIME SERIES. Each of the nine panels shows the modeled binary range surface for one temporal bin across the study period, derived by thresholding the ensemble climatic suitability surface. Panels are arranged chronologically from left to right and top to bottom. Predicted presence range is shown in color; predicted absence in gray. The black outline delineates the USGS GAP range boundary for the species.

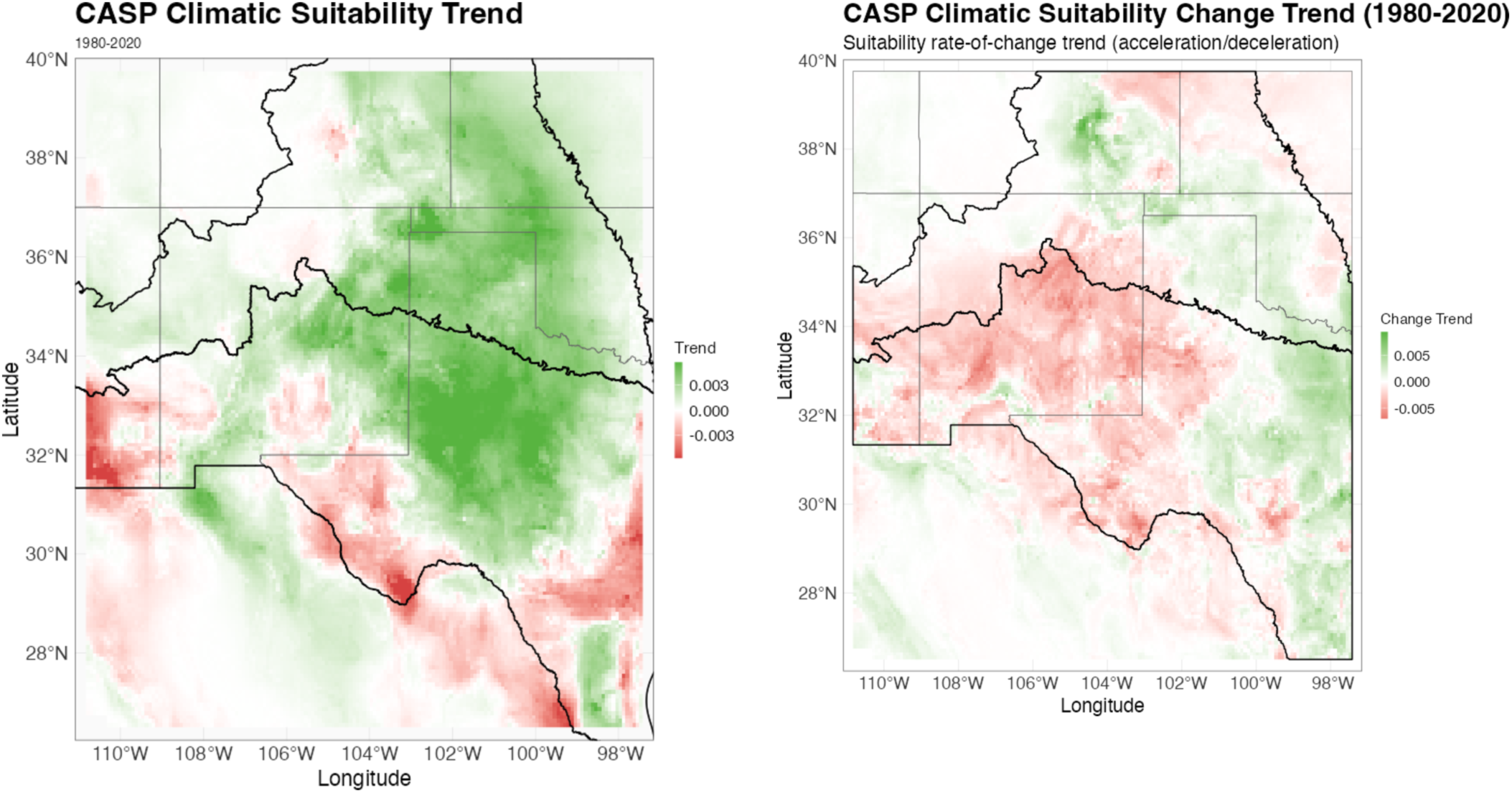

CLIMATIC SUITABILITY TRENDS. The left panel shows the per-cell Theil-Sen trend in climatic suitability across the modeled extent over the study period. Green indicates increasing suitability; red indicates decreasing suitability. The right panel shows the trend in the rate of suitability change over the same period, identifying areas where change is accelerating or decelerating. The black outline delineates the USGS GAP range boundary for the species.

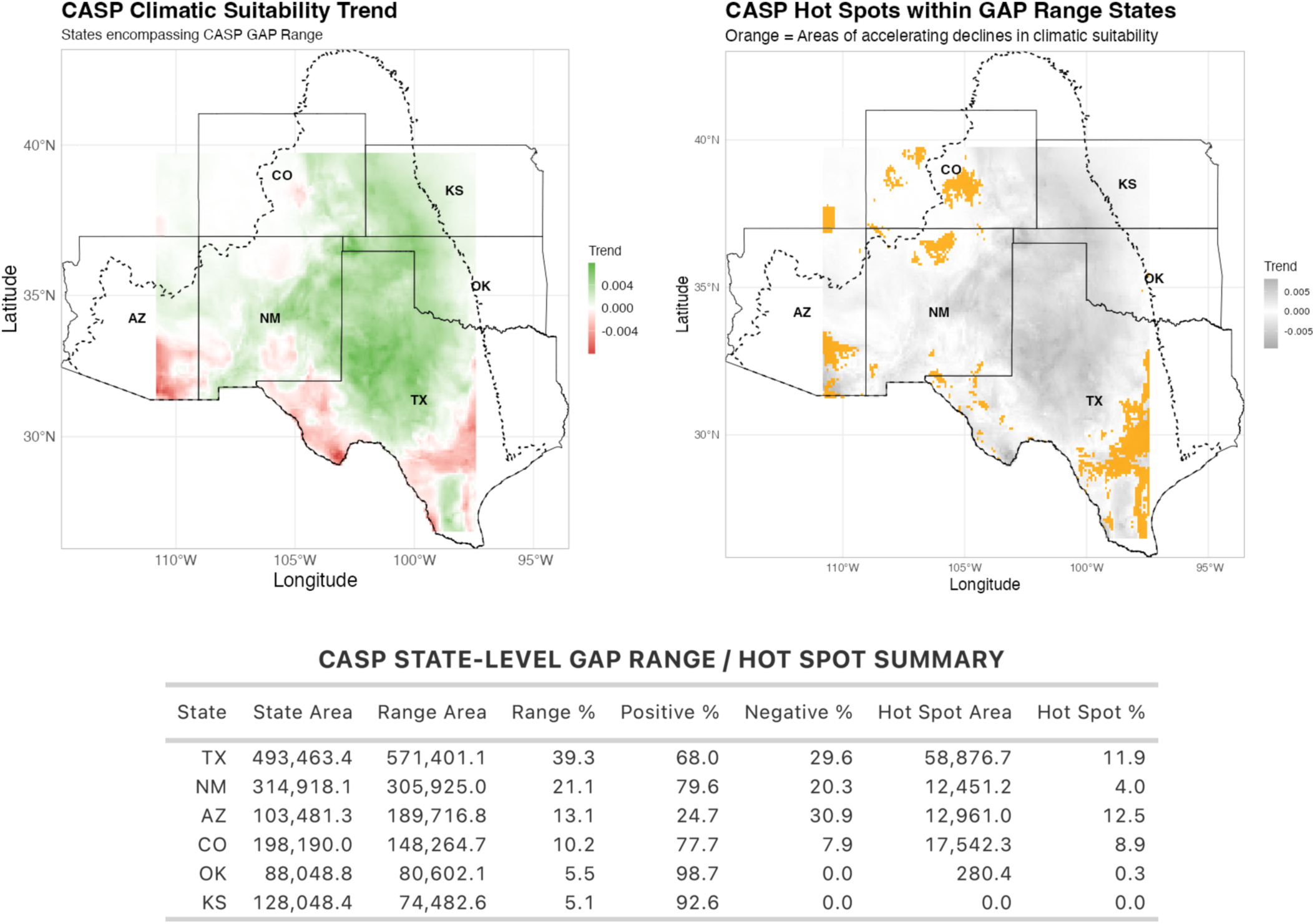

STATE-LEVEL SUITABILITY TREND AND HOTSPOT SUMMARY. The left map shows the per-cell Theil-Sen suitability trend restricted to states encompassing the highest percentages of the species’ USGS GAP range. Green indicates increasing suitability; red indicates decreasing suitability. The right map shows hotspots of accelerating suitability decline within those same states; orange cells are areas where the rate of suitability loss is itself increasing. The table reports, for each state, total state area, GAP range area and percentage, the proportions of range cells with positive and negative trends, hotspot area, and hotspot percentage of range area. All areas in km^2^.

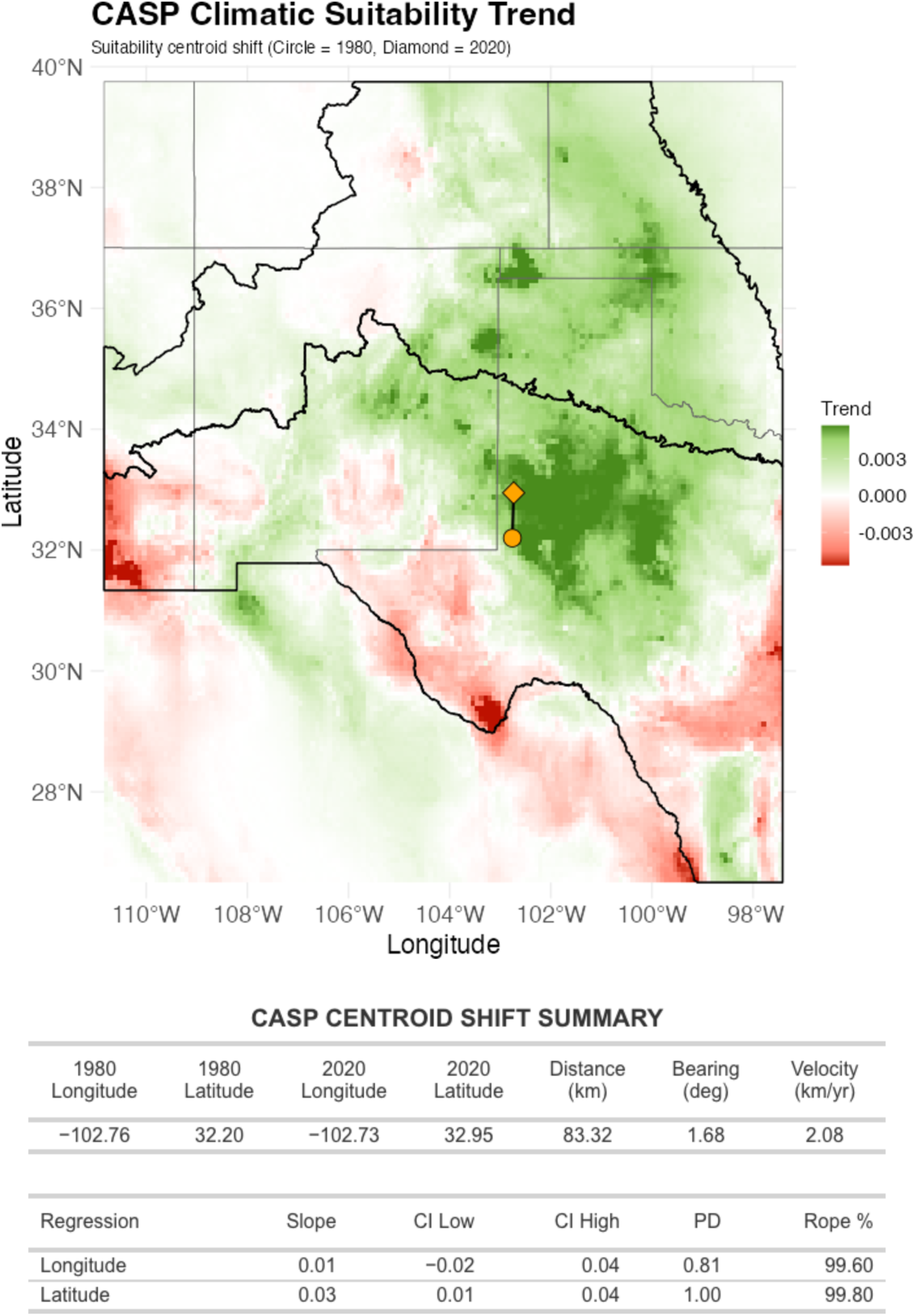

CLIMATIC SUITABILITY TREND WITH CENTROID SHIFT. The map shows the per-cell Theil-Sen trend in climatic suitability across the modeled extent over the study period. Green indicates increasing suitability; red indicates decreasing suitability. The circle marks the suitability-weighted centroid in the first temporal bin; the diamond marks its position in the last temporal bin. The black outline delineates the USGS GAP range boundary. The table reports centroid displacement distance, bearing, and velocity over the study period, with Bayesian regression statistics for the longitudinal and latitudinal centroid trajectories across temporal bins.

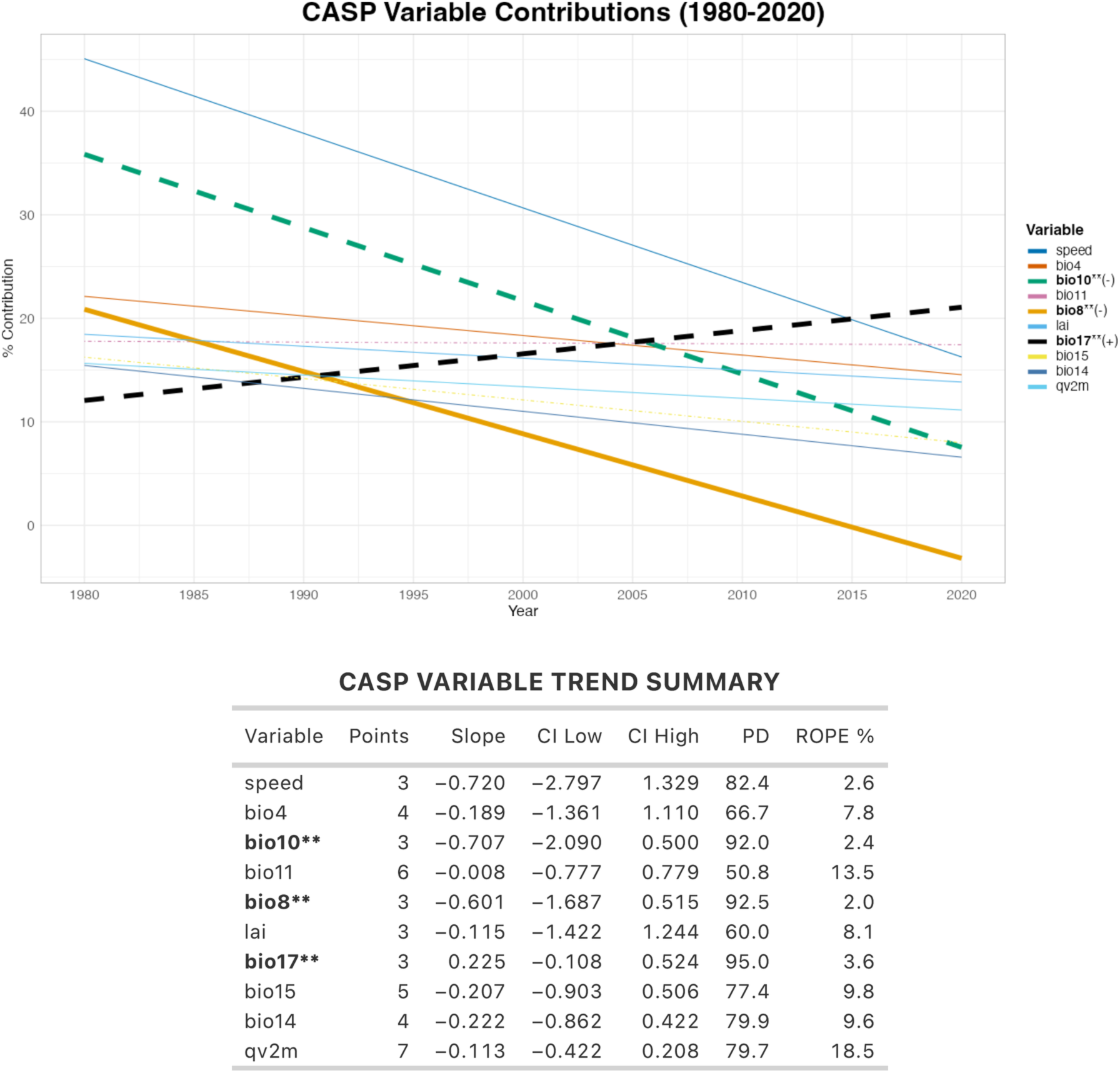

VARIABLE CONTRIBUTION TRENDS. The plot shows how the relative contribution(% contribution) of each top environmental predictor to the ensemble models changes across temporal bins over the study period. Each line is a Bayesian regression fit to one variable’s contribution trajectory; the legend indicates the direction of that trend where statistically supported(+ increasing, -decreasing). The table reports, for each variable, the number of temporal bins in which it appeared among the top predictors (Points), the Bayesian regression slope, 95% credible interval bounds (Cl Low, Cl High), probability of direction (PD), and percentage of the posterior within the region of practical equivalence (ROPE %). Variables in bold show strong directional trends.

PREDICTOR VARIABLE TRENDS. The following pages show how the top contributing MERRA-2 and MERRAclim-2 environmental predictor variables have changed across the landscape over the study period. Variables are those identified by the convergence-based screening procedure as the strongest contributors to the ensemble models; they are arranged in descending order of average contribution to the overall time series.

Each variable occupies one row. The left panel shows the per-cell Theil-Sen trend over the study period. The right panel shows the trend in the rate of change over the same period, identifying areas where the variable is increasing or decreasing at an accelerating or decelerating rate. In both panels, green indicates increasing values and red indicates decreasing values.

A blue sidebar marks variables whose relative contribution to the ensemble models showed a strong directional trend across the study period (Bayesian probability of direction PD ≥ 90%). See the Variable Contribution Trends figure and table for details.

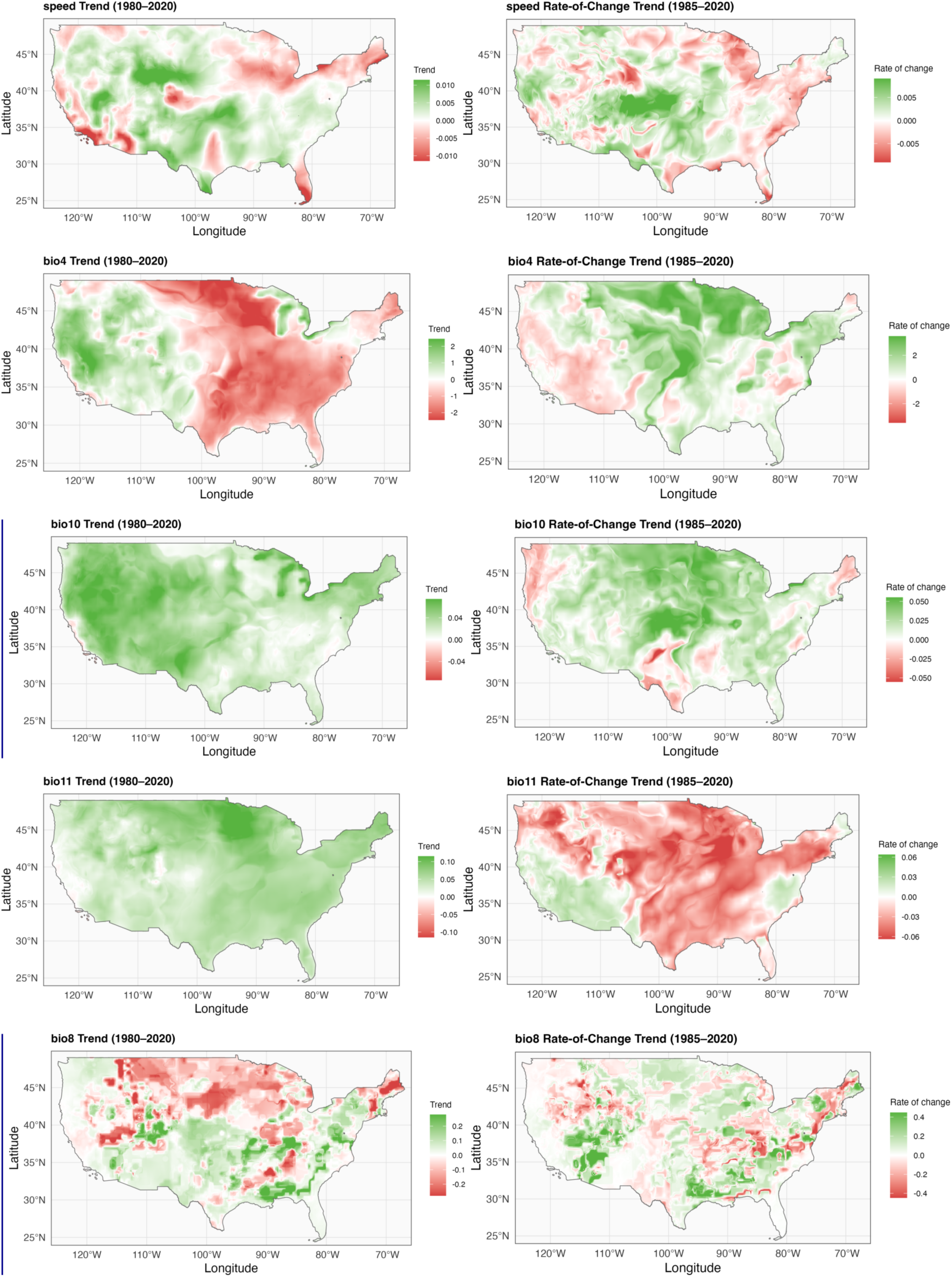

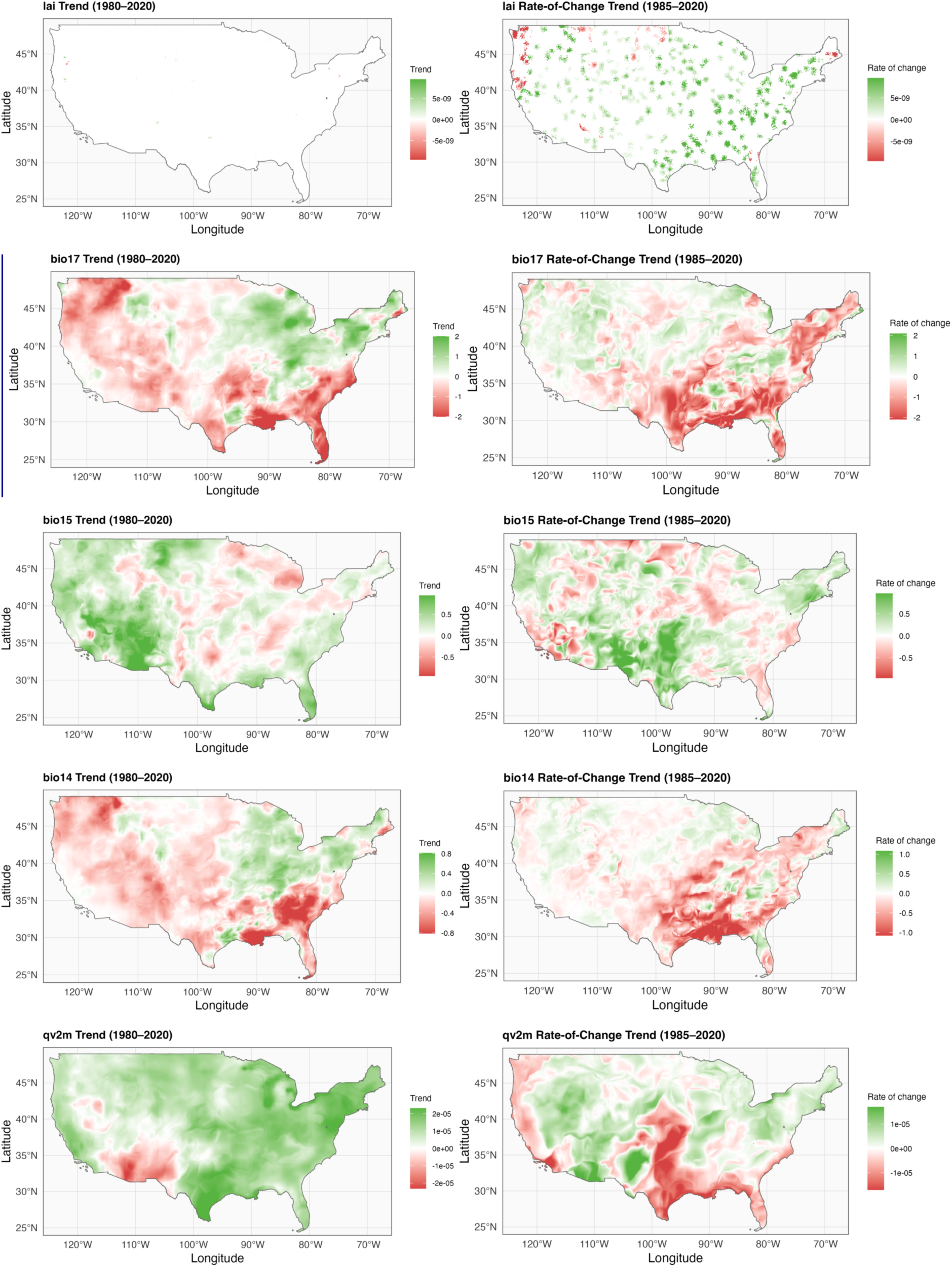

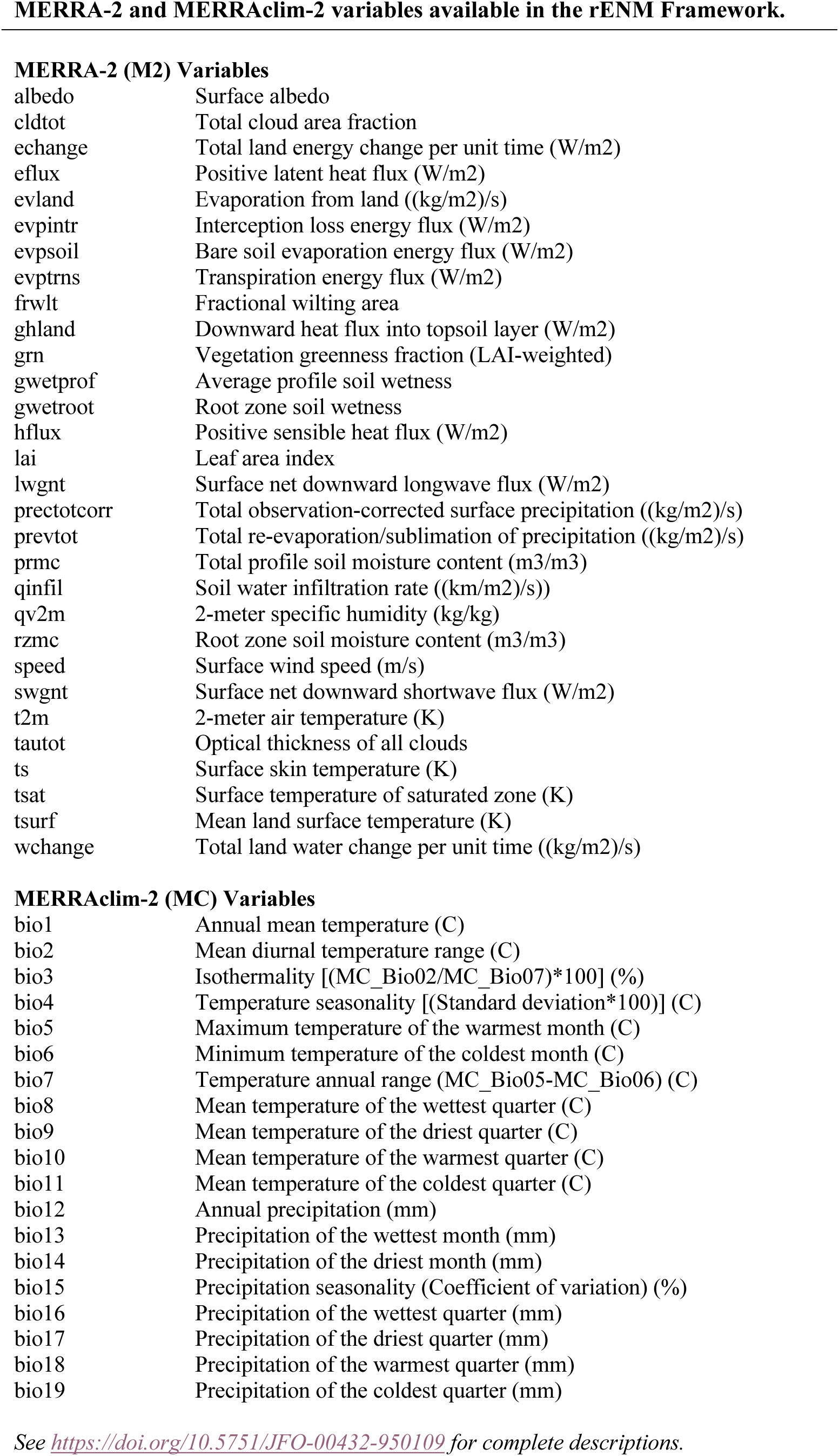

